# Ripples of Resistance: Unveiling Antimicrobial Resistance Dynamics Along Switzerland’s Aare River

**DOI:** 10.64898/2025.12.18.695068

**Authors:** Denise Lea Wälchli, Sasikaladevi Rathinavelu, Jane Ackeret, Norberto Aquino, Karin Beck, David J Janssen, Helmut Bürgmann

## Abstract

The global spread of antimicrobial resistance (AMR) is a serious public health concern, driven by widespread antibiotic use and the global environmental circulation of antibiotic-resistant bacteria and resistance genes (ARGs). Wastewater treatment plants (WWTPs) are important sources of anthropogenic AMR entering large rivers, which serve as vital water resources but facilitate downstream dissemination. The drivers and dynamics of AMR propagation along river systems remain poorly understood. As Switzerland’s longest and one of its largest rivers, the Aare, situated in the upper Rhine watershed, plays a central role in the ‘water castle of Europe’. This study examines the impact of WWTP discharges, some receiving high loads of hospital effluent, on ARG distribution along the 288 km Aare river-continuum. Using quantitative PCR targeting 14 ARGs conferring resistance to eight antibiotic classes, combined with *16S rRNA* gene amplicon sequencing, we conducted a high-resolution spatial survey to assess shifts in the riverine ARG content and microbiome. Concentrations of trace metals and nutrients were analyzed as tracers of anthropogenic inputs. Results revealed a progressive increase in ARG abundance downstream, driven by WWTP effluents enriched in ARGs. Effluents had 70-fold higher mean ARG concentrations than upstream waters, raising downstream levels up to 141-fold. Major tributaries such as the Reuss and Limmat sustained elevated ARG levels, while passage through lakes markedly reduced concentrations. This study provides the first detailed baseline for ARG prevalence along a large river system, from pristine headwaters to pollution-affected lower reaches and insights into aquatic AMR dynamics and guidance for future monitoring.

## 1. Introduction

Bacterial antimicrobial resistance (AMR) has emerged as a major global health and development threat of the 21^th^ century^1^. In 2021, an estimated 4.71 million deaths were associated with AMR, including 1.14 million deaths directly attributable to bacterial AMR^1^. According to projections, by 2050 AMR could be responsible for 8.22 million deaths, underscoring the urgent need for global action^1^.

Although antimicrobial resistance genes (ARGs) occur in pristine environments, the levels of resistance observed today are not solely due to naturally occurring resistances, e.g. in antibiotic-producing bacteria^2–5^. Instead, the widespread use of antimicrobials in healthcare and livestock farming has significantly contributed to the spread of antibiotic-resistant bacteria (ARB) and ARGs in ecosystems^6–8^. Human and animal wastes contaminated with antibiotics, ARB and ARGs are continuously released to the environment, either untreated (e.g., agricultural run-off, aquaculture, sewer overflows) or partially treated in wastewater treatment plants (WWTPs). Generally, WWTPs receiving wastewater (WW) from various anthropogenic sources of AMR, including urban areas, hospitals, and specific industries (e.g., pharmaceutical industry and industrial slaughterhouses) are not designed for efficient micropollutant removal or disinfection^9–13^. Consequently, WWTPs have been found to be a major source of ARGs, facilitating their dissemination into aquatic environments^10,14^. While WWTPs quantitatively reduce ARG and ARB loads^15,16^, they also serve as hotspots for ARG persistence, horizontal gene transfer (HGT), and bacterial adaption, with effluents exhibiting high transcriptional activity and highly mobilized ARGs^16–18^.

Aquatic ecosystems may then contribute to ARG dissemination by acting as a reservoir and conduit of received resistance determinants. This enables acquisition, evolution, and recombination of resistance genes through HGT of mobile genetic elements (MGEs) in pathogenic and clinically relevant bacteria, driving the emergence of new resistance traits^19^. Effluent discharge has well-documented impacts on the resistome of downstream river ecosystems, with ARG primarily persisting in the water column^20,21^. While ARG levels were observed to decrease over short distances (∼2 km) from a point source by dilution, and potentially also via sedimentation and degradation, mass balance and fluctuations observed over longer distances (> 10 km) suggest AMR removal, local proliferation and influence of non-point sources on riverine AMR levels^22^. This highlights the role of aquatic environments in the long-term spread and evolution of AMR.

Rivers serve as critical interfaces between humans, animals, and the environment. Flowing for hundreds of kilometres, they act as conduits for WW discharge, carrying (micro)pollutants (e.g., organic chemicals, pharmaceuticals, pesticides, and detergents), as well as ARB and ARGs to downstream ecosystems across landscapes and even continents^9,10,23^. While antibiotics are generally expected to degrade, although at variable rates, the case is less clear for ARB and ARGs^24^. These can decrease through various mechanisms but can potentially also persist for decades^25^ or even proliferate under suitable conditions^22^. These contaminants can re-enter the microbiomes of human and animal populations when river water is used for drinking, irrigation, and recreational activities, potentially leading to colonization or in the worst case, causing skin, gastrointestinal, urogenital, and respiratory infections with ARB^26–28^.

This interconnectedness is of particular importance in Switzerland, often called the “Wasserschloss Europas” (engl. “Water castle of Europe”) due to its extensive river networks and vital freshwater resources feeding into major European river systems such as the Rhine, Rhone, and Danube. Swiss rivers shape regional water quality and contribute to the broader European hydrological system, affecting pollution levels in waters across the Swiss border^29^. Protecting these waterways is essential for mitigating the spread of AMR, safeguarding drinking water sources, and maintaining the ecological and public health of large parts of Europe.

Previous studies on AMR in larger river systems exposed to strong anthropogenic pressure have mostly targeted heavily polluted sites. While recent research has focused on large European rivers like the Danube^30^ and Rhine^31^, as well as rivers in China^32–35^ and the USA^21,36^, no comparable studies have been conducted in Switzerland. In general, studies with high-resolution longitudinal sampling and simultaneous examination of potential point sources of contamination are lacking.

This study aimed to establish the anthropogenic impact on AMR prevalence along Switzerland’s Aare River, from its source to its confluence with the Rhine River. Originating from the Ober-and Unteraar glaciers in the pristine alpine Grimsel region and thus lacking upstream pollution, it flows downstream through increasingly agricultural areas and densely populated cities, where anthropogenic impact increases. We focused on a quantitative approach, relying on quantification of AMR indicator genes using qPCR and included ARG concentration, relative abundance and mass-flow perspectives. We explored the hypothesis that WWTP effluents in general, and those from WWTP receiving hospital WW in particular, are a main source of AMR that largely explains the observed accumulation. Additionally, the study explored potential sinks along the river course, assessing whether AMR is removed, persists, or proliferates. Finally, we investigated whether this system’s microbial community structure drives AMR dynamics.

## 2. Materials and Methods

### 2.1. Study area and sample collection

The Aare, Switzerland’s longest river, drains approximately 43% of the country (catchment size: 17709 km^2^) with a mean annual discharge of 555 m^3^/s (Gauge Station Siggenthal^37^). The Aare River originates in the high Alps, fed by glacial sources, and flows over 288 km through landscapes characterized by increasing intensity of agricultural and industrial use, population density and urbanization prior to its confluence with the Rhine (**Fig. 1a**). **S1.1** provides a detailed description of the river. Switzerland’s temperate climate, with year-round precipitation and distinct seasonal variations, strongly influences discharge, with spring snowmelt being a major contributor.

**Figure 1:**
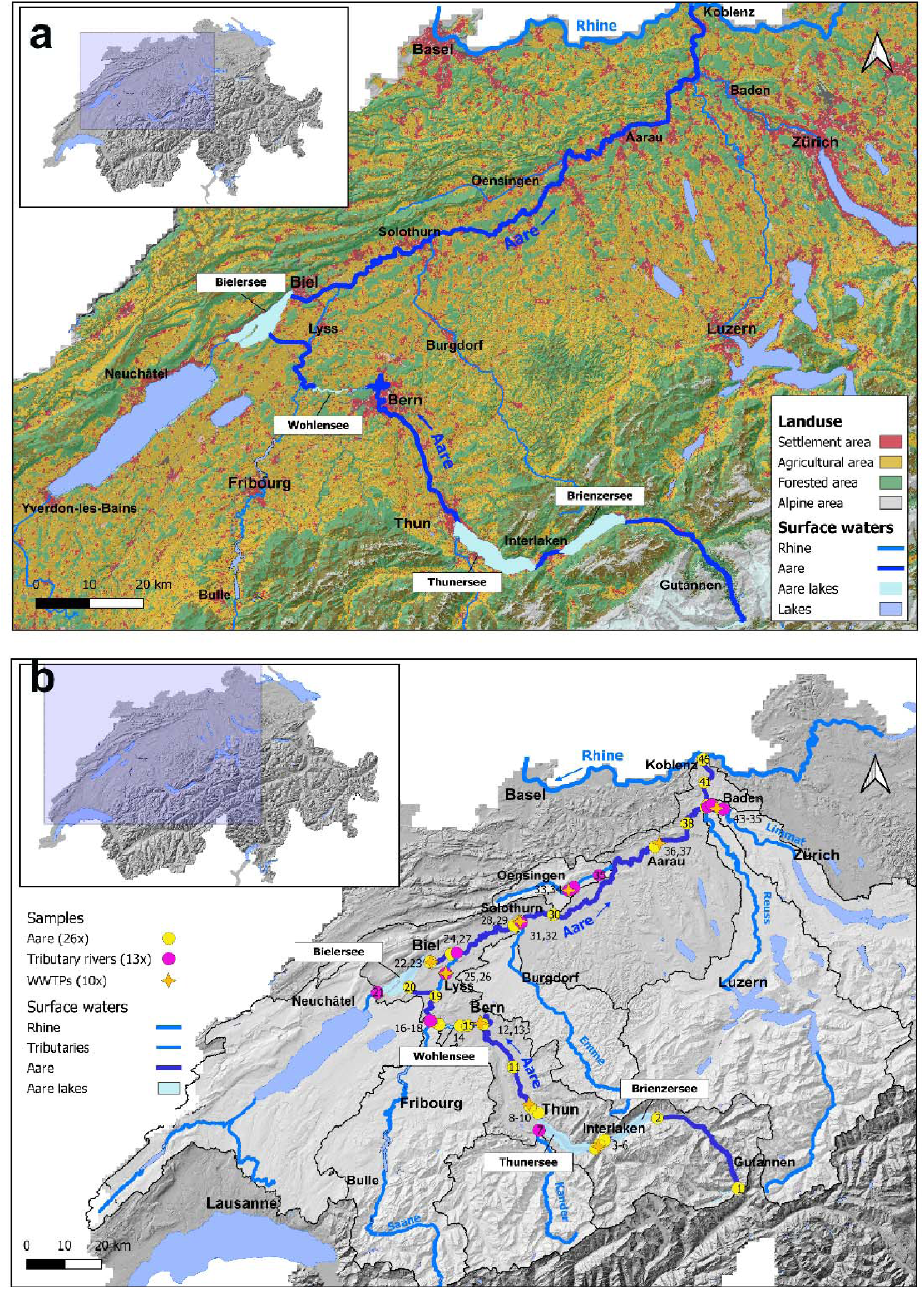
Land use and sampling sites within the Aare River catchment, Switzerland. (a) land-use distribution in the Aare catchment. High-altitude unproductive alpine areas are shown in grey tones, forested areas in green, agricultural areas in yellow, and settlements in red. (b) Map illustrating 49 sampling locations in the Aare River, its tributaries, and from selected municipal WWTPs receiving WW from either industrial slaughterhouses (2 samples) or hospitals (8 samples). Sampling point numbers are indicated (details in Table S1). The map indicates topography (relief). The Aare catchment is highlighted and tributary river sub-catchment borders are shown as black outlines. In both panels, lakes are shown in blue, and lakes traversed by the Aare River (dark blue) are shown in light blue. Blue arrows indicate the direction of flow of the Aare and Rhine Rivers.

The field campaign (04 to 14 March 2024) targeted expected low flow conditions prior to snowmelt, thus conditions in which the impact of wastewater would be near the maximum. In total, 49 samples were collected from the Aare River, its main tributaries and potential contamination sources (**Fig. 1b, Tab. S1**). Effluents of nine municipal WWTP discharging into the Aare River and its tributaries were sampled, seven of which also serve large hospitals (Interlaken, Thun, Bern, Biel, Solothurn, Aarau, and Baden). Two WWTPs (Lyss and Oensingen) receive pre-treated WW from industrial slaughterhouses. At Lyss, an additional sample was collected at the pre-cleaning step. Six tributaries with catchments > 500 km^2^ (decreasing catchment size: Reuss > Zihl > Limmat > Saane > Kander > Emme) were sampled near their confluence with the Aare River. Samples downstream of WWTP or tributary inflows were taken 6 – 10 km downstream, where possible, to ensure complete mixing. For samples from the Aare, and points of WWTP discharge and tributary confluence, the distance from the source (in river kilometers, rkm) was determined from GPS coordinates of sampling points and a shapefile of the Aare River in QGIS Maidenhead^38^.

Samples were primarily collected from bridges to capture mid-river water at ∼30 cm depth. A Comet-Combi 12 4T pump (Comet Pumps, Pfefferschwanden, DE) was used. To avoid contamination, the pump, tubing and 5 L plastic sample bottles were rinsed thrice by pumping ∼ 5-10 L of river water prior to sampling. Per location 2 x 5 L samples were taken over ∼5 minutes for biological analyses except for WWTP effluents (1 x 5 L). Subsamples were obtained for microbial, chemical, and nutrient analysis. A separate grab sample was obtained with a metal-free sampling bottle for trace metal analyses. Metadata (temperature, pH, conductivity, and dissolved oxygen) were recorded in situ using a WTW multi-parameter device, MultiLine 3630 IDS (Fisher Scientific, USA).

### 2.2 Flow cytometry (FCM)

Total cell count (TCC), and ratios of high nucleic acid (HNA) and low nucleic acid (LNA) cells were determined by FCM on a NovoCyte Advanteon flow cytometer (Agilent, USA) according to the Swiss accredited Standard method 333.1, based on SYBRgreen staining^39^. The procedure followed the methodology described by Proctor and Besmer et al.^40^. Negative controls with sterile-filtered Evian water were included. The gating strategy was recalibrated to ensure accurate differentiation of microbial cells from other particles (see **S1.2, Fig. S1**).

### 2.3 Biomass Filtration in the Field

Aliquots of all collected water samples were filtered in the field using S-Pak mixed esters of cellulose Membrane filters (GSWG047S6, Millipore, USA) with a pore size of 0.22 µm and a diameter of 47 mm. Filters were placed in sterilized polysulfone filter holders (Ref. No. 10461100) with a diameter of 50 mm (Whatman, Germany) under sterile conditions. A peristaltic pump type V6-3L (Shenchen, China) with two pump heads equipped with silicon tubing (# 35 from the manufacturer) was utilized for the filtration process. From each sample, two filter replicates were produced and stored immediately at-20 °C until DNA extraction. Filtration volume (**Tab. S1**) depended on the turbidity of water samples. To avoid contamination between samples, silicon tubing was rinsed with 70% ethanol, followed by nanopure water.

### 2.4 DNA Extraction

DNA was extracted from both filter replicates using the DNeasy PowerWater Kit (Qiagen, Germany) following the manufacturer’s protocol. To optimize yield, the elution volume was reduced to 65 µL per replicate, and the extracts were subsequently pooled, resulting in a final DNA sample volume of 130 µL. DNA concentration was quantified using the Qubit 4 Fluorometer (Invitrogen, USA), and purity was assessed with the NanoDrop One spectrophotometer (Thermo Fisher Scientific, USA) (see **Tab. S2**). Blank extractions confirmed that filters and reagents were free of DNA contamination.

### 2.5 Quantitative PCR (qPCR)

The abundance of a total of fourteen ARGs as indicators for anthropogenic AMR input (*aadA, aph(3’)-lb*, *bla_CTX-M-1_, bla_TEM_*, *ermB*, *ermF*, *mcr-1*, *mecA*, *sul1*, *sul2*, *tetA*, *tetM*, *tetW*, *qnrA)* was determined by qPCR. In addition, qPCR was performed for four genetic markers (GM): class 1 integron integrase gene *intI1* as a marker for mobile (multi-)resistance cassettes; CrAssphage (crAss) as a marker for tracking human fecal contamination^41^; the beta-glucuronidase gene *uidA* as an *E. coli* specific marker for fecal indicator bacteria, and bacterial *16S rRNA* genes (*16S*) as a proxy for bacterial abundance^42,43^. qPCR assays were performed on a LightCycler®480 (Roche, Switzerland). Standard dilutions (5 x 10^7^ to 50 copies/reaction) and negative controls (*no*-*template*-control: AE buffer; PCR-control: only PCR reagents without the addition of sample) were run in technical quintuplicates, while 1:10 and 1:100 dilutions of each pooled extract and extraction blanks were analyzed in triplicate. Plasmid and synthetic DNA standards were used for calibration. Primer sequences, primer concentration, amount of DNA, and cycling conditions are provided in **S1.3** and **Tab. S3 – S7**. Information on quantification standards is provided in **Tab. S8-S9**. Gene abundances were obtained by normalizing to sample volume, logarithmized (base 10) and reported in units of log copies/L. Calculations for LOD, LOQ, qPCR efficiency, and PCR data analysis, as well as the relative abundance of anthropogenic AMR indicators normalized to *16S* followed the methodology outlined by Rathinavelu et al. and Czekalski et al.^44,45^. qPCR assays met qualitative standards for linearity (R^2^ > 0.99) and efficiency (90-110%), with the exception of the *tetA* gene (R^2^ = 0.981) (see **Tab. S7**).

For each WWTP location, we calculated AMR contamination potential (F_EFF/US_) and observed WWTP impact (*F_DS/US_*) as log-transformed ratios of gene abundance (*C*) in effluent (*C_EFF_*) or downstream (*C_DS_*) samples, respectively, to the upstream river samples (*C_US_*) (**Eq. 1, 2**).

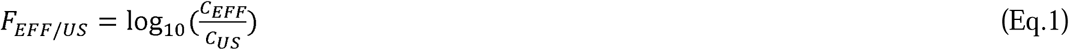

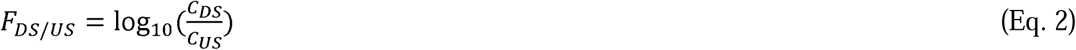

### 2.5 *16S rRNA* Gene Amplicon Sequencing

Microbial community composition was assessed by *16S rRNA* gene amplicon sequencing of the V3-V4 region (**S1.4**), performed with PCR-cDNA Barcoding Kit SQK-PCB111.24 on GridION platform using MinION flow cells R9.4.1 (Oxford Nanopore Technologies, UK). Sequencing depth was a minimum of 30,000 reads per sample. Quality control included re-sequencing of samples that showed matrix inhibition. Resulting FASTQ data were matched against the SILVA database (Release 138, Ref Nr. 99) using EMU^46^. Non-bacterial sequences were removed prior to data analysis.

### 2.6 Metal, Chemical, and Nutrient Parameters

Total nitrogen (TN) and total organic carbon (TOC) were determined following ISO standards (TN: ISO 11905-1:1998, TOC: ISO 1484:1979) using a Skalar San++ system (Procon AG, Switzerland) and a Shimadzu TOC-L analyzer (Japan). Anions (fluoride, chloride, sulfate) were determined by anion chromatography (930 Compact IC Flex, Metrohm, Switzerland). Dissolved trace metals (Copper (Cu), Zinc (Zn) and Gadolinium (Gd)) and Phosphorus (as total dissolved Phosphorus (TDP) and total Phosphorus (TP) in filtered and unfiltered samples respectively) were quantified by Agilent 8900 ICP-MS equipped with a PrepFAST autosampler, following a protocol adapted from Janssen et al. (2024)^47^. Raw data were processed with MassHunter 5.1 (Agilent). Measurements were conducted in four batches (Sept. 2024 – Apr. 2025) with procedural blanks and a certified reference material (SLRS-6) in each batch (**Tab. S10**). Additional details are provided in **S1.5**.

### 2.7 Hydrological Data and Total ARG Load

To evaluate discharge dynamics during the campaign weeks from source to estuary, discharge data from nine gauging stations (**Tab. S11**) along the Aare River were obtained from the Swiss Federal Office for the Environment^48^. These data were used to calculate mean daily discharge for each gauge (**Tab. S11**) and then to model sampling point discharge by interpolating between gauging stations and accounting for discharge of major tributaries (**Tab. S11, Fig. S2)**. We then calculated ARG loads (*L*) for each sampling point in the Aare River and each ARG by multiplying discharge values (*Q_i_*) at each sampling location (*i*) with ARG abundances according to **Eq. 3**.

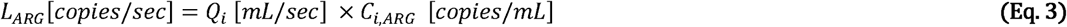

Additional discharge and load calculations were conducted for sampling locations in four major tributaries (Saane (rkm 134.8), Emme (rkm 196.5), Reuss (rkm 272.6), and Limmat (rkm 274.1)) using data from the gauging station closest to the confluence, and for WWTP effluent discharge using data provided by the WWTP (**Tab. S10, S11**). To assess if WWTP discharge explains observed ARG increases downstream of sampled WWTP, we compared observations to predicted ARG gene concentrations downstream of WWTP (*C_DS_*) obtained from a simple discharge-based mixing model (**Eq. 4)**.

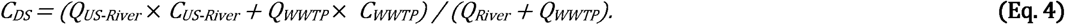

Where *Q_WWTP_*is measured WWTP discharge, *Q_US-River_* is modeled river discharge and *C_US-River_ and C_WWTP_* are measured ARG concentrations in the upstream river (US) and WWTP effluent (EFF).

### 2.8 Data Processing and Visualization

Data handling, statistical analyses and graphics were generated in R^49^ using R Studio^50^ and the packages ggplot2, cowplot, stringr, scales, grid, readr, knitr, dplyr and tidyr. Species richness and Shannon’s alpha diversity index were calculated using the vegan^51^ package and differences among groups were tested using the Kruskal-Wallis’ test provided in base R. Non-metric multidimensional scaling (NMDS) was performed on *16S rRNA* abundance data with Bray-Curtis distance as the dissimilarity measure (k = 4). Differences in microbial community composition between different waterbodies were tested using PERMANOVA. Spearman rank correlations were performed to assess relationships between microbiological, chemical, nutrient parameters, and concentrations of trace metals. QGIS Maidenhead^38^ was used for geographical illustration and map design. Adobe Photoshop was used to fine-tune graphical illustrations.

## 3. Results

### 3.1 Background Characterisation of Anthropogenic Influence on the Aare River

Nutrient and metal concentrations reflect anthropogenic impact of urban and agricultural activity along the Aare River. Both TDP (total dissolved phosphorus), representing the bioavailable, short-term reactive fraction, and TP (total phosphorus), reflecting total P-load, remained < 10 µg/L upstream of rkm 118, but concentrations increased further downstream (**Fig. 2c**). WWTP effluents emerged as major point sources, with TP >100 µg/L (except WWTP Thun, 60 µg/L), while tributary P concentrations were generally similar to the Aare River at confluence.

**Figure 2:**
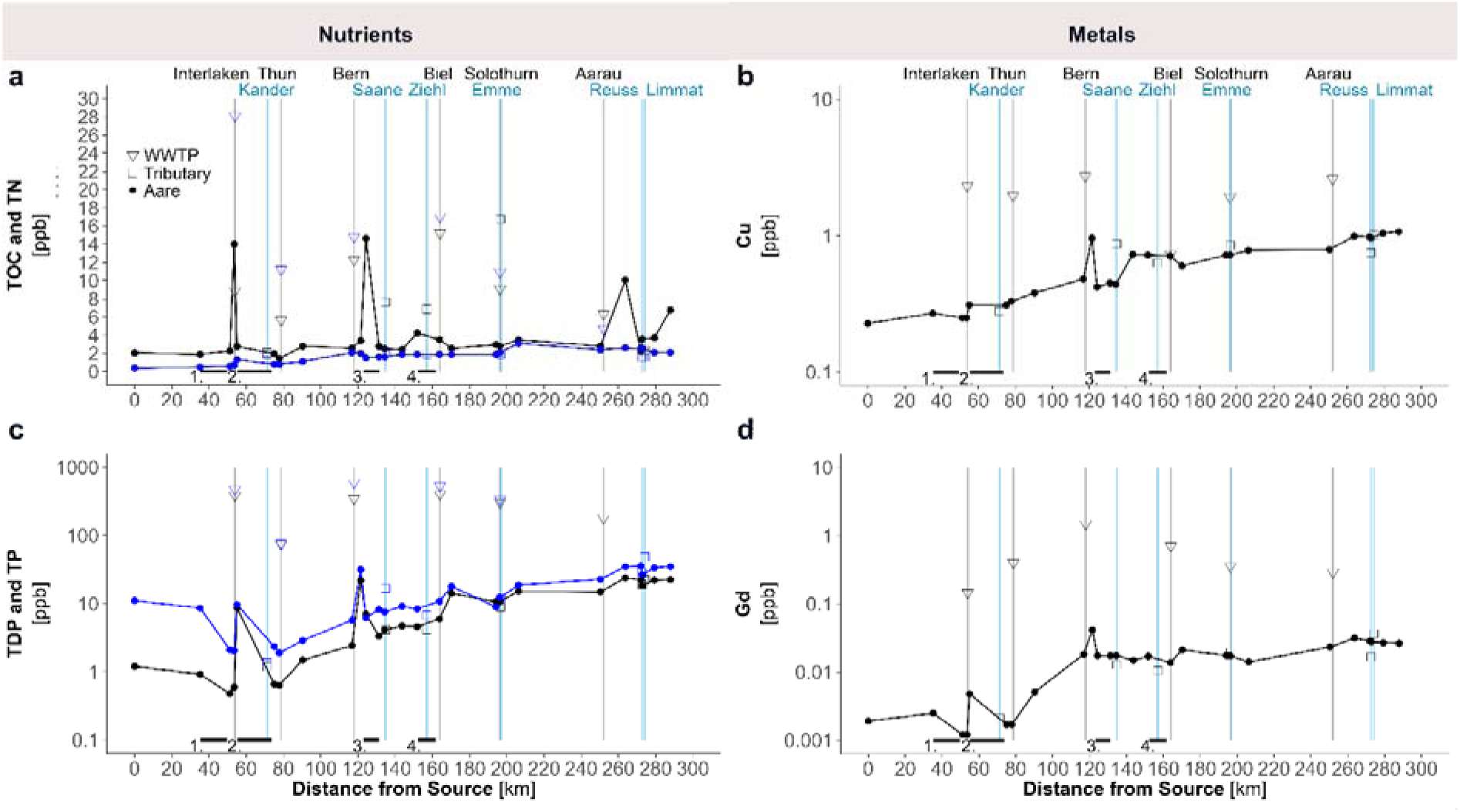
Dynamics of nutrients and metals along the Aare River. Left-side panels show nutrient dynamics: (a) total organic carbon (TOC; black) and total nitrogen (TN; blue), and (c) total dissolved phosphorus (TDP; black) and total phosphorus (TP; blue). Right-sided panels display (b) copper concentrations and (d) gadolinium concentrations. In all panels, values along the river course are shown as continuous lines, while values from WWTP effluents (triangles on grey bars) and tributaries (squares on blue bars) represent input values. Flow-through lakes are indicated as black bars on the x-axis: 1) Brienzersee, 2) Thunersee, 3) Wohlensee, and 4) Bielersee.

Three trace metals were compared to further characterize anthropogenic impacts: Copper (Cu) and Zinc (Zn) as indicators of diffuse urban, agricultural, and industrial activity^52,53^ and gadolinium (Gd) as a specific tracer of healthcare-related inputs^54^, reflecting its use as contrast agent in magnetic resonance imaging (MRI). Copper concentrations gradually increased from 0.22 µg/L at the source to 1.07 µg/L at the confluence with the Rhine. WWTPs were point sources with up to 2.6 µg/L Cu (**Fig. 2b**). Zn followed a similar pattern (**Fig. S3**), but in contrast to Cu also increased in the downstream section between Solothurn and Aarau, consistent with non-point sources such as road runoff and industrial emissions^52^. Gd concentrations showed a similar trend, but with more pronounced increases downstream of WWTPs receiving hospital effluent, notably Interlaken (0.0048 µg/L), Bern (0.04 µg/L), and Aarau (0.03 µg/L). WWTP effluent Gd concentrations were > 0.1 µg/L, 10^1^ – 10^2^ fold higher than river background. Furthermore, elevated Gd/Cu ratios, especially in Thun, Bern, Biel and Solothurn WWTP effluents, reflect the specific point source input of Gd through WWTP of urban regions with healthcare centres relative to the more varied Cu sources, confirming Gd as a sensitive tracer of healthcare-related discharges (**Fig. S4**).

### 3.2 Trends in Microbial Community Abundance and Composition

The observed trends in nutrient and metal distribution are consistent with microbial indicators such as TCC and high HNA fraction, consistent with elevated microbial abundance at nutrient-enriched sites (**Fig. 3a, Fig. S5**). TCC values steadily increased roughly ten-fold from the source to the Rhine, ranging from 1.7 x 10^5^ to 1.7 x 10^6^ cells/mL. Tributaries draining urban areas (e.g., Saane, Reuss, Limmat) showed elevated TCCs, whereas those from more pristine regions (e.g., Emme, Kander) were lower than the Aare at their confluences. *16S rRNA* gene sequencing further revealed spatial patterns in microbial diversity and composition (**Fig. 3b, 3c**). Beta diversity analysis using NMDS on Bray-Curtis dissimilarities demonstrated structuring by spatial proximity, with headwater (rkm 0-35) and upper Aare (rkm 35-100) communities being significantly different from more tightly clustered downstream communities (rkm >100) (PERMANOVA: R² = 0.323, F = 1.813, p = 0.001, **Fig. 3b**). Tributaries fell into the cluster of the river stretch they entered, except Zihl channel, while composition of WWTP effluent communities was clearly differentiated.

**Figure 3:**
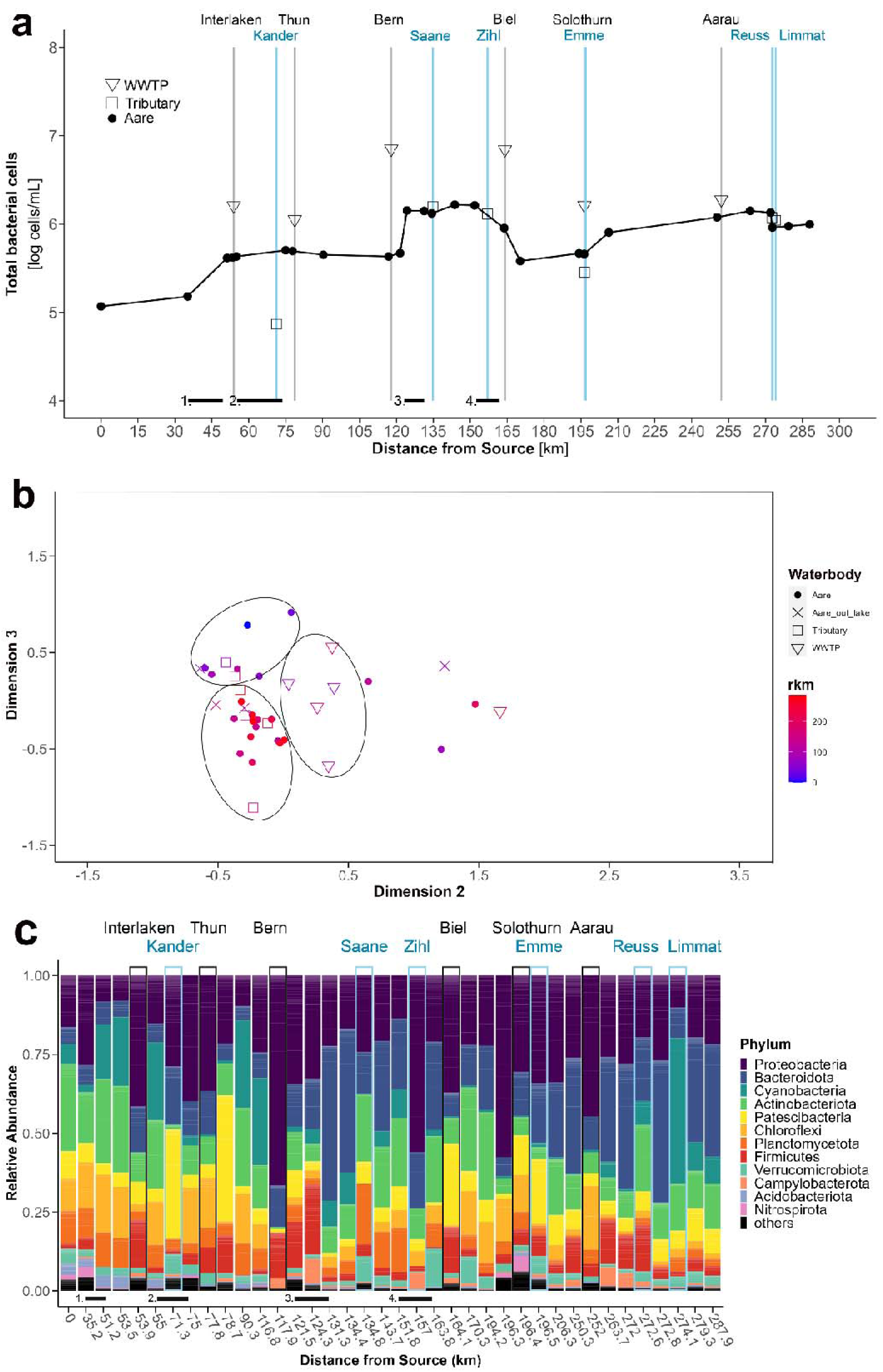
Composition and dynamics of the bacterial community along the Aare River. (a) Total bacterial cell count (TCC) along the river course (black line), including effluents from WWTPs along Aare River (triangles on grey bars), and tributaries (square on blue bars). (b) NMDS ordination showing sample distribution along the Aare River based on community composition, highlighting three aggregates (black circles): i) WWTP effluents, ii) upper reach samples, and iii) lower reach samples. (c) Relative abundance of the 12 most frequent bacterial phyla along the Aare River, its tributaries (blue bars), and WWTP effluents (black bars); remaining phyla are grouped as “others”. Flow-through lakes are indicated as black bars on the x-axis: 1) Brienzersee, 2) Thunersee, 3) Wohlensee, and 4) Bielersee.

These trends in community composition are clearly visible in the phylum level taxon distribution (**Fig. 3c**). Localized shifts in the community were noted e.g., downstream of Thunersee (increase in *Patescibacteria*), around Bielersee (increased *Cyanobacteria*) and in metropolitan areas downstream of WWTPs such as Bern and Aarau, where a notable rise in the abundance of *Firmicutes* and *Campylobacterota* was observed. In WWTP effluents, *Proteobacteria* were consistently dominating, and together with *Bacteroidota* comprising up to 70% of the sequences. *Patescibacteria* were particularly abundant (> 25%) in the WWTP effluents of Thun and Biel and *Firmicutes* were enriched (> 10%) in effluents of WWTPs Interlaken, Thun, Bern, and Biel. In the Aare and its tributaries, considerable variability was observed.

### 3.3 Unique ARG Distribution Patterns Along the Aare River

Out of 14 investigated ARGs, 11 (*aadA*, *aph(3’)-lb*, *bla_CTX-M-1_*, *bla_TEM_*, *ermB*, *ermF*, *sul1*, *sul2*, *tetA*, *tetM* and *tetW*) were detected in the Aare River (**Fig. 4a**). *qnrA*, *mecA*, and *mcr-1*, which confer resistance to quinolones, methicillin, and colistin, respectively, were not detected in any of the Aare River samples, but *qnrA* was found in two WWTP effluents (**Fig. 4d**). Some ARGs, e.g. those conferring resistance to aminoglycosides (*aadA*), erythromycin (*ermB*), sulfonamides (*sul1*), and tetracycline (*tetW*) were frequently detected and typically the most abundant (**Fig. 4a, Tab. S1**). In contrast, β-lactamase resistance genes (*bla*_CTX-M-1_, *bla*_TEM_) and the *E. coli* marker gene *uidA* were restricted to low-level detections (< 10^2^ copies/mL) at hotspots with high overall ARG concentration (rkm 124 and 250).

**Figure 4:**
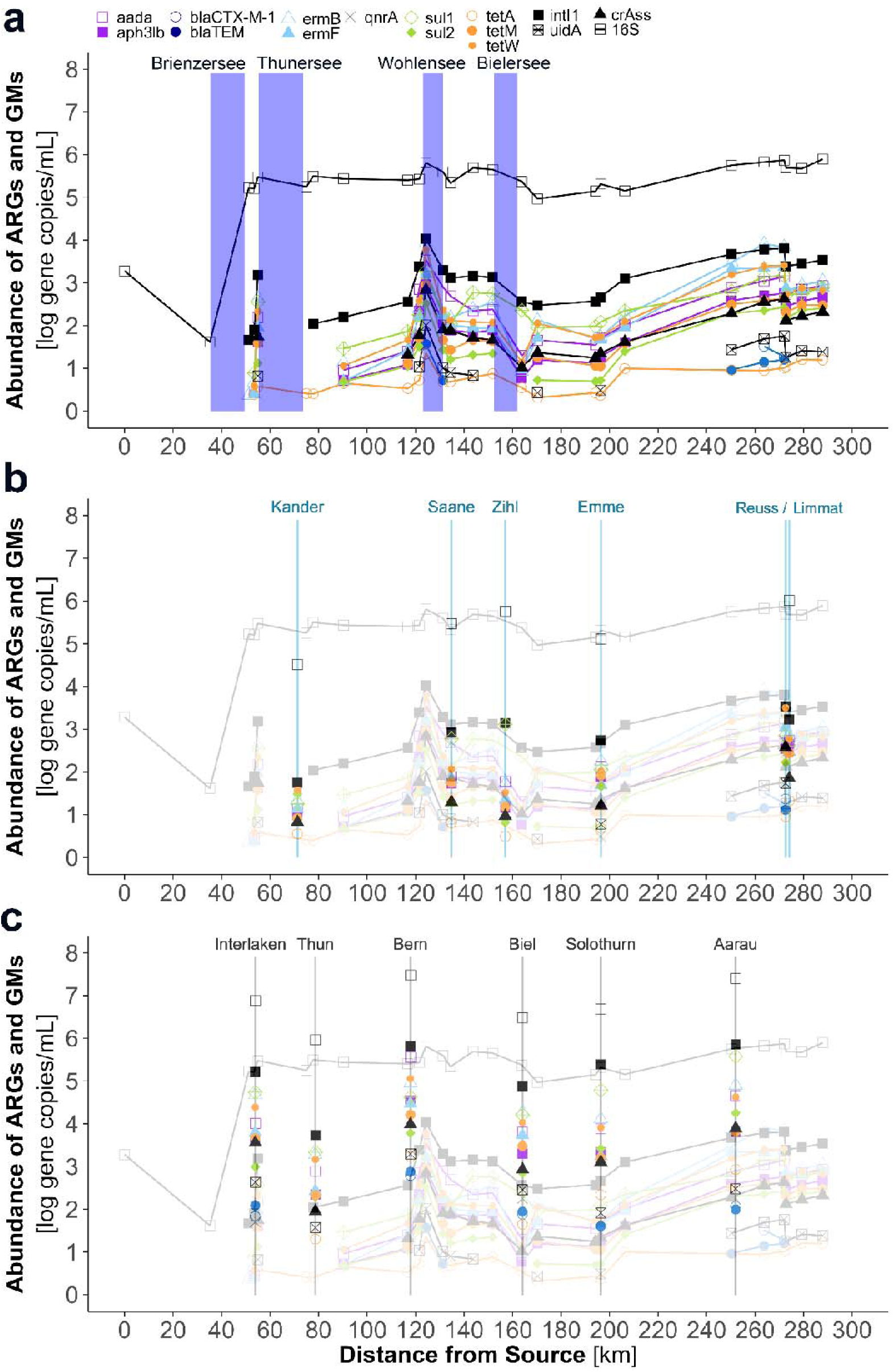
Absolute abundance (log gene copies/mL) of ARGs and GMs, quantified using qPCR. (a) The concentration trends for ARG and GM in the Aare River, with the location of flow-through lakes indicated as blue bars. The ARG abundances measured in (c) tributary inflows and (d) WWTP effluents is depicted as symbols within vertical lines indicating the inflow or confluence location (tributaries: blue / WWTP: gray), superimposed over the Aare River data from panel (a) (faded lines). The values are logarithmic, - 1 corresponds to 10 and 2 corresponds to 100 copies/mL.

At the source of Aare River and at rkm 30, ARGs were not detected, while *16S* was present at ∼1.9 x 10^3^ copies/mL (**Fig. 4a**). From rkm 45 onwards, *16S* copies increased, consistently exceeding 1 million copies/mL close to the confluence. ARGs and the integron integrase gene *intI1* were first detected at the outflow of Brienzersee near Interlaken, although at relatively low concentrations (10^1^ – 10^3^ copies/mL) and were detected downstream. *intI1* was highly prevalent along the entire downstream river (up to 10^4^ copies/mL) and, with few exceptions, the most abundant AMR indicator. Both absolute (**Fig. 4a**) and relative ARG abundances (**Fig. S6a**) as well as their total load (i.e. fluxes; **Fig. S7**) increased overall along the river’s course. The increase was non-continuous, with local declines in ARG abundance at rkms 75, 130, 165 and marked peaks or increases at rkms 55, 124, 170, 206 until reaching a maximum between rkm 250 to 272 (**Fig. 4a**) with ARG diversity broadening in the lower reaches (detections of *bla*_CTX-M-1_, *bla*_TEM_).

Total load calculations showed that the flux of the *16S* marker gene increased almost 70-fold (5.84 x 10^12^ to 4.06 x 10^14^ copies/sec) while *intI1* increased 1090-fold (1.63 x 10^9^ to 1.78 x 10^12^ copies/sec) between rkm 51 (ARG detected) and rkm 288 near the confluence. *ermB* increased 7120-fold (8.1 x 10^7^ to 5.77 x 10^11^ copies/sec), with similar trends for other widespread ARGs, indicating the river’s role in downstream ARG transport (**Fig. S7**, **Tab. S12**).

### 3.4 ARG Patterns of Aare River Shaped by Lakes, Large Tributaries, and WWTPs

#### Lakes

When ARGs were present upstream of lakes, a pattern of “resetting” ARG abundance was observed when the river passed through a lake (**Fig. 4a**). Thunersee eliminated all ARGs, which had reached a first peak downstream of the city of Interlaken, to levels below detection, with the exception of *tetA*. In contrast, Wohlensee and Bielersee caused a drop in ARG abundance but did not eliminate previously detected ARGs after lake passage, except for *tetM* and *sul2*, which were undetectable for ∼7 rkm downstream of the Bielersee outflow. Although no measurements upstream of Lake Neuchâtel were performed, the low ARG abundance in the Zihl canal connecting Lac de Neuchâtel and Bielersee (**Fig. 4c**) is also consistent with this pattern.

#### Tributaries

Across all sampled tributaries, the same ARGs (*aadA*, *aph(3’)-lb*, bla_CTX-M-1_, *bla*_TEM_, *ermB*, *ermF*, *sul1*, *sul2*, *tetA*, *tetM,* and *tetW*) and GMs (*16S*, *intI1,* and *uidA*) were detected as in the Aare River (**Fig. 4b**). Meanwhile, ARGs *qnrA*, *mecA*, and *mcr-1* remained undetected in all tributaries. ARG abundance was positively correlated with population density within the catchments (Spearman ρ = 0.89, p = 0.033; **Fig. S8**), showing tributaries draining larger and more populated areas exhibit higher ARG abundances. Among the tributaries with smaller catchment areas (< 2000 km^2^), the Kander exhibited consistently low concentrations of all markers (< 100 copies/mL), consistent with low population density. In comparison, Saane (1893 km^2^) and Emme (976 km^2^) rivers, draining more populous regions, showed elevated levels of ARGs, reaching up to 1000 copies/mL The Limmat River (2412 km^2^) draining the Zurich metropolitan region showed an even higher abundance of ARGs, including *tetA*, which was detected at even higher levels than in the Reuss river (3426 km^2^). The Reuss drains major urban regions in the Cantons Lucerne and Zug, and large parts of central Switzerland, and is the tributary with the largest catchment and overall highest ARG load. At the confluences with Reuss and Limmat, Aare River displayed higher concentrations of both ARGs and GMs compared to the inflowing tributaries, which resulted in a dilution effect downstream of the confluence (**Fig. 4b**).

#### Wastewater

WWTP effluents consistently contained high ARG abundances exceeding those in river water (**Fig. 4c**). In addition to *uidA* and *intI1*, 9 out of 14 ARGs were detected across all effluents, including genes conferring resistance to aminoglycosides (*aadA*, *aph(3’)-lb*, sulfanomides (*sul1*, *sul2*), tetracyclines (*tetA*, *tetM*, *tetW*), and macrolides (*ermB*, *ermF*). *bla_TEM_* was absent only in WWTP Thun, while *bla*_CTX-M-1_ was found only in effluents from WWTPs Interlaken, Bern, Biel, and Aarau. *qnrA* appeared at low levels (50 – 180 copies/mL) in WWTP Interlaken and Biel. *mcr-1* and *mecA* were not detected. Downstream of WWTPs, most notably for Interlaken (rkm 54) and Bern (rkm 118) an increase in ARG levels in the Aare River was observed and analyzed in further detail.

The observed contamination downstream of WWTP (log DS/US ARG abundance ratio; F_EFF/US_, **Fig. 5a**) was consistent with the WWTP contamination potential (log EFF/US ARG abundance ratio, F_DS/US_, **Fig. 5b**). Inflows of WWTP effluents, with on average 70-fold (geometric mean; average F_DS/US_: 1.9) higher ARG concentrations than upstream river water (F_DS/US_ range:-0.2 to 4.2, **Fig. 5b**) led to ∼4-fold (0.6 log) downstream increases of ARGs in the receiving river (F_DS/US_ range:-0.4 to 2.1, **Fig. 5a**). Negative F_DS/US_, indicating ARG abundance at or below background levels, were only observed for the effluents of the pre-clean stage of the slaughterhouse at Lyss and the smallest WWTP, Baden.

**Figure 5:**
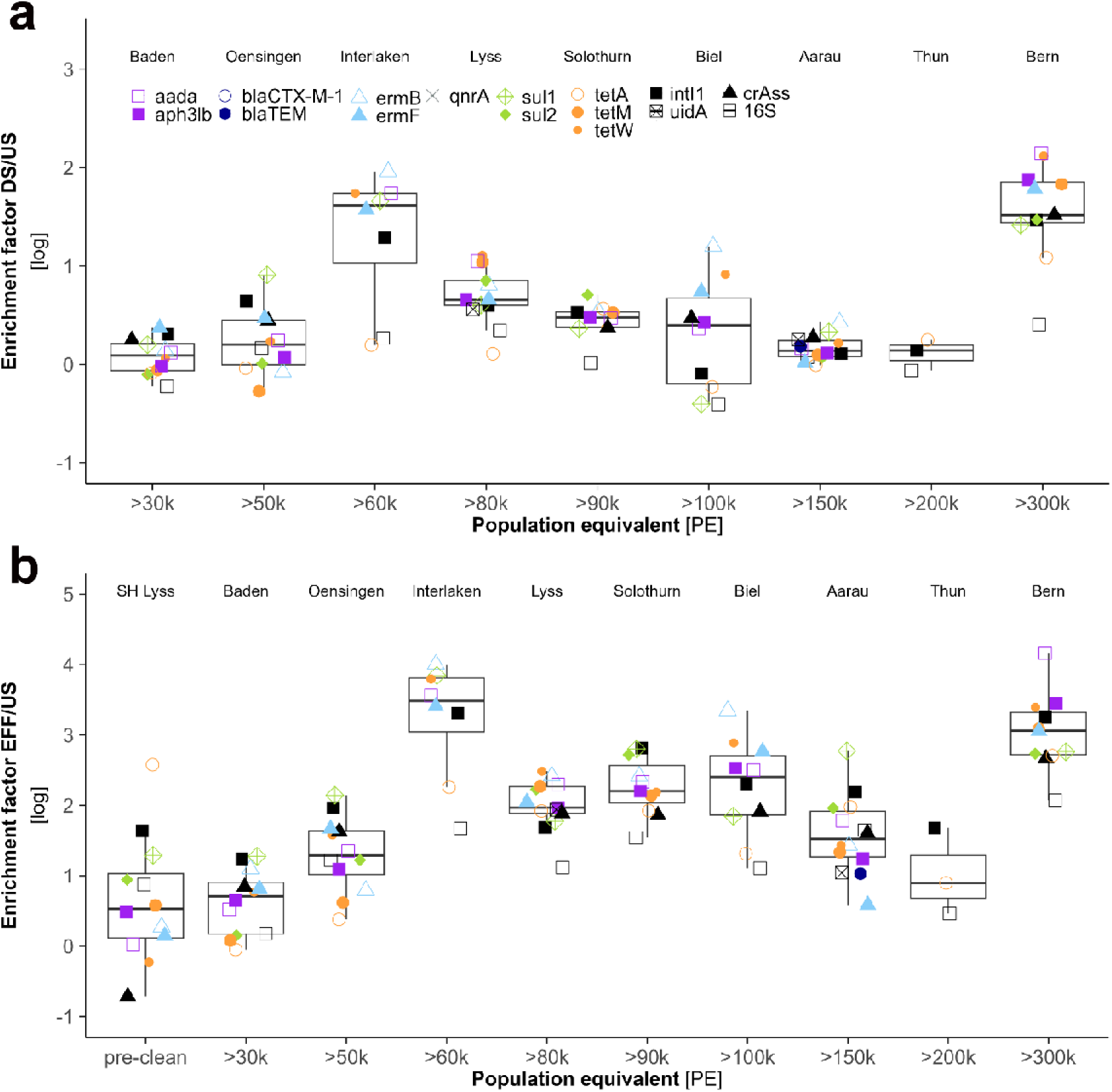
Observed contamination (DS to US ratio) and contamination potential (EFF/US ratio) of the absolute abundance of ARGs and GMs with increasing design capacity (PE) of WWTPs. For evaluation of contamination potential and the impact of WWTP effluent on ARG and GM abundance in receiving river, log10-transformed ratios of ARG abundance were calculated for (a) downstream relative to upstream, and (b) WWTP effluent relative to the upstream river. Results were ordered by the design capacity (population equivalent, PE) of WWTPs. Three WWTPs located on tributaries of the Aare River were also included: Baden (Limmat), Lyss (Alte Aare), and Oensingen (Dünnern). The latter two receive wastewater from slaughterhouses in addition to communal wastewater.

The contamination potential (log EFF/US ratio, F_EFF/US_) of the effluent was similar for all genes, the Kruskal-Wallis test (p = 0.489) and pairwise Wilcoxon comparisons found no statistically significant difference of F_EFF/US_ between genes across WWTP (**Fig. 5b**). The large WWTP in Bern (design capacity > 300 kPE), which also receives WW from the Inselspital (University Hospital) had the highest ARG contamination potential with a median F_EFF/US_ of 3.1. The small WWTP of Lyss (pre-clean, <30 kPE, F_EFF/US_ =0.53) and Baden (<50 kPE, F_EFF/US_ =0.71) also had the lowest ARG contamination potential. However, no significant relationship (R^2^ = 0.14, p = 0.29) was observed between ARG contamination potential and design capacity overall (**Fig. S9**).

For F_DS/US_, Kruskal-Wallis test (p = 0.223) and Wilcoxon pairwise test again showed no significant difference between genes. The F_DS/US_ for crAss was also not statistically distinct from ARG, indicating that both can be explained by WWTP impact. WWTP impact varied by site (**Fig. 5a**). The strongest median contamination impact (1.66) was observed at WWTP Bern (> 300 kPE). The increase was particularly high for *aadA* (141-fold), *aph(3’)-lb* (75-fold), *tetW* (132-fold), *tetM* (68-fold), and *ermF* (61-fold). No impact (i.e. negative F_DS/US_) was observed more frequently (n=14) than for contamination potential, mainly at WWTP Baden, Oensingen and Biel.

Applying a discharge-based mixing model, the expected downstream ARG concentration matched observations closely (R^2^ = 0.88, p = 0.005, m=0.80, **Fig. S10**), indicating that the observed increases are largely explained by the impact of the WWTP effluent, without taking additional sources or non-conservative behavior of AMR into account. Previous work has shown that ARG typically behave similar to conservative chemical tracers over distances of a few km although other processes have an impact over greater distances^22^. The impact of WWTP of urban centers thus consistently resulted in a marked increase in ARG concentrations, loading, and relative abundance in the Aare River. This is most pronounced downstream of WWTP Interlaken and Bern (**Fig. 4a, d**) but is also contributing to the cumulative increases downstream of Biel (rkm 164-170), Solothurn (rkm 196-206), and up to Aarau (rkm 206-272).

An analysis of the relationship between ARG relative abundance and microbial community composition in Aare river samples (S1.4, S3) indicated common environmental gradients drive community and ARG structure. Procrustes analysis revealed a significant concordance between microbial community composition and ARG profiles (r = 0.47, p = 0.003). dbRDA showed that a subset of ARGs (*aada, aph3lb, sul1*) best explained independent variation in community composition, which is in line with anthropogenic impacts, such as wastewater discharge, driving both community composition and ARG content (see S3).

### 3.6 Correlation between ARG and other indicators of anthropogenic impact

To identify factors associated with variations in the abundance of ARGs and GM (*intI1*, *16S,* crAss and *uidA*), we calculated Spearman rank correlations with microbiological and chemical parameters and concentrations of nutrient and trace metals. Correlation analyses were performed separately on samples collected i) along the Aare River (**Fig. 6**) and ii) on effluents from 10 WWTPs along the Aare River and its tributaries (**Fig. S11**). To capture spatial trends along the river continuum, “km from source” (distance) was included as a parameter.

**Figure 6:**
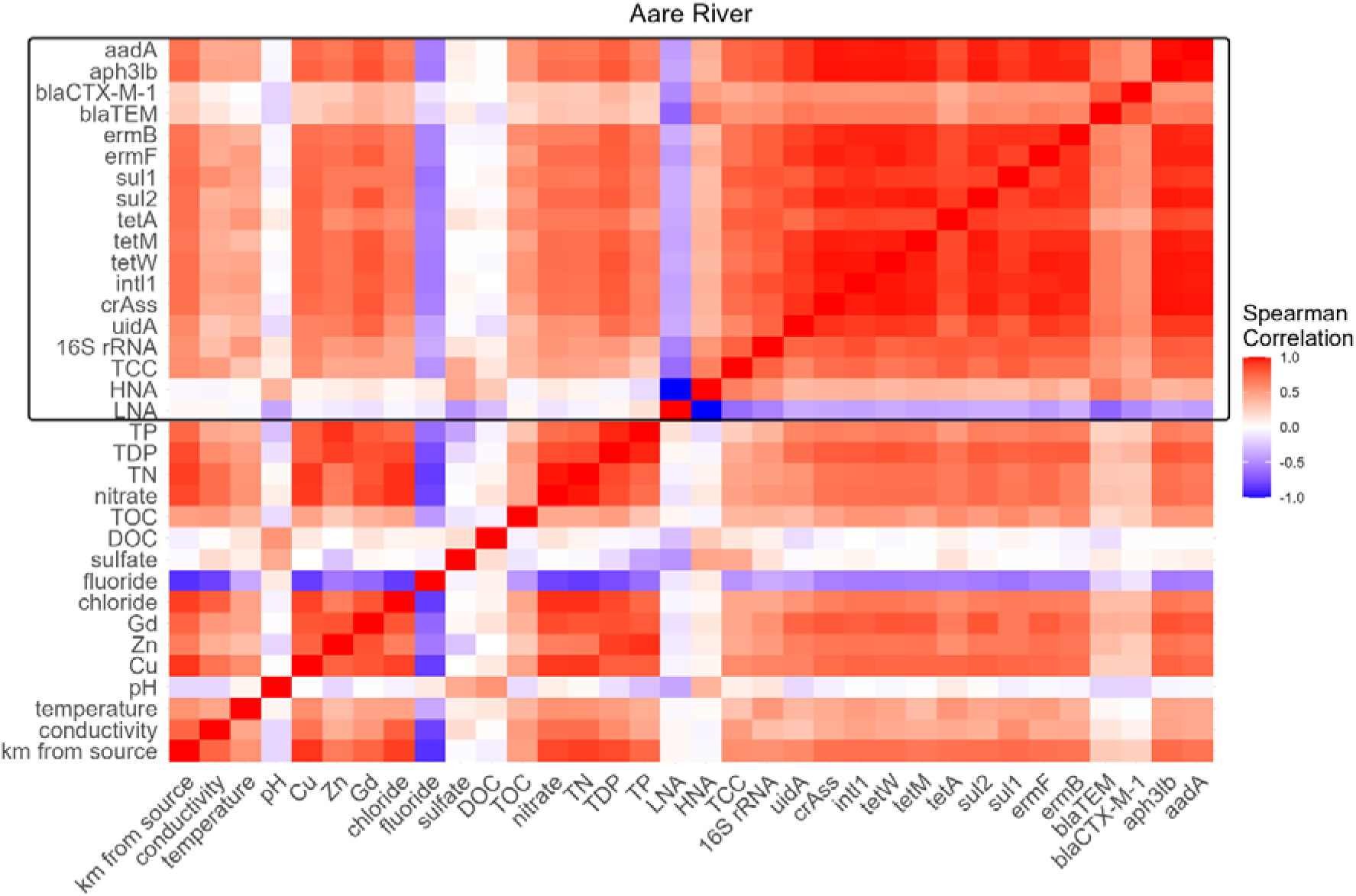
Heatmap of Spearman correlation matrix including quantified genes, physicochemical parameters, and biological indicators across all Aare River samples. Color indicates the spearman correlation coefficient (ρ). The black box indicates biological parameters, including ARGs, GMs, and FCM-derived cell counts.

In river water samples, we observed positive correlations between most parameters, with the exception of fluoride, sulfate, pH, DOC, and LNA abundance (**Fig. 6**). Distance emerged as a strong predictor of most parameters, including ARG abundance. Strong positive correlations (ρ > 0.7) were observed between distance and the abundance of *aph(3’)-lb*, *ermB*, *intI1*, *sul1*, *sul2*, *tetA*, and *tetW* and the human fecal source tracking indicator crAss. But also nutrient parameters like TP, TDP, chloride, nitrate, and TN (ρ > 0.8) and trace metals, particularly Cu (ρ = 0.92) increased with distance.

Across the dataset, most ARGs showed highest correlation coefficients with one another (ρ > 0.85), except for both β-lactamase genes (*bla_CTX-M-1_*and *bla*_TEM)_, which displayed only moderate correlations with other ARGs (ρ = 0.4–0.7) due to values below detection in many samples, but co-occurred strongly with each other (ρ > 0.8). ARGs also correlated closely with *intI1* indicating that ARGs mobilized into class1 integron cassettes likely contributed to ARG dynamics. Typically class-1 integron associated genes like *sul1* and *aadA* were highly correlated. *16S, TCC,* and *uidA,* but especially crAss abundances correlated with ARG occurrence (ρ > 0.85, omitting β-lactamase genes), supporting their roles as indicators of total bacterial load and faecal contamination, and a mostly human source of ARG. Notably, Gd correlated highly with nearly all ARGs (ρ > 0.65), with the rare β-lactamases being the only exception.

Correlation analysis of WWTP effluent samples revealed very strong correlations among most ARGs, including *bla_CTX-M-1_* and *bla*_TEM_, while *tetA* consistently exhibited weak correlations (**Fig. S11**). ARGs correlated mostly positively with fecal contamination marker crAss and *uidA*, nutrients, metals, and bacterial abundance measures, but less closely (mostly ρ < 0.5) compared to river water samples. Distance was not a strong or consistent predictor for ARGs or other parameters in the WWTP effluent dataset.

## 4. Discussion

### 4.1 WWTPs are major contributors to the ARG content of the Aare river

Overall, observations on the River Aare were consistent with urban sources and WWTP being the key contributors to ARG accumulation. For all WWTP sites studied, effluent was shown to be a source of ARG relative to the receiving river water (**Fig. 5a, b**), which accounts for most of the downstream ARG increase observed near major urban centres (**Fig. S10**). This remains the case even in the lower reaches of the Aare, e.g. at Aarau, where upstream contamination has accumulated considerably. Accumulation of ARG also clearly exceeded the total bacteria (2.5 median DS/US enrichment of *16S*) showing that ARG enrichment represents an effect distinct from total bacterial input. Close correlation of ARG and crAssphage abundance supported a primarily human (fecal) origin of ARG in the Aare river. Our campaign was conducted pre-snowmelt, and despite some precipitation at near-minimum discharge^55^, water temperature and UV-radiation ^56^. With dilution and removal mechanisms like predation, starvation-, and UV-induced mortality probably being at a minimum, our data likely represents a near-maximum impact.

Along the river continuum, two main hotspots of locally increasing ARG content at Bern (rkm 118) and Interlaken (rkm 54) highlighted how source and sink processes can drastically affect downstream dynamics. In both cases, sharp peaks in ARG absolute and relative abundance downstream of the WWTP were short-lived, as the Aare entered a lake just after each city (**Fig. 4a, c Fig. S6b, d**). The most pronounced effect was observed in Bern, which was also the largest WWTP sampled. On an average, effluent showed a 3.1 log higher ARG content compared to upstream water, resulting in the highest downstream increase (1.6 log). WWTP Interlaken, despite operating at only one-fifth the capacity of Bern, showed an even higher mean effluent ARG content relative to upstream water (3.2 log). While partly explained by the comparatively pristine upstream water, Interlaken’s role as a tourism hotspot may have contributed to higher absolute abundances of ARG compared to e.g. Thun (**Fig. 4d, 5b**). According to the local tourist association, more than 3.5 million overnight stays per year add substantially to the local population (https://www.interlaken.ch/info-service/ueber-interlaken-tourismus/berichte-statistiken/jahresberichte), with WW from visitors, residents, and the regional hospital all processed by the WWTP.

Previous work has shown that many ARG in Swiss wastewater are frequently associated with mobile genetic elements ^16^ including conjugative plasmids^45^ and part of multiresistance-conferring genetic systems^57^, indicating an elevated hazard potential and the potential to accelerate resistance evolution through gene transfer. The observed high abundance of *intI1* and its correlation with ARG in the Aare are in line with this expectation, but we did not study the role of mobile genetic elements in more detail in this study.

Our results support earlier observations on the importance of WWTP discharge for ARG levels in smaller Swiss rivers with proportionally high wastewater impact^22^, and demonstrate this also for a major river system along its full length.

### 4.2 ARG as a component of anthropogenic impact – common sources and selection

Other markers of human activity such as P and Cu (with sources in agriculture and urban wastewater) increased in parallel with the AMR markers (**Fig. 2, 4**). The increase in Gd (MRI contrast agent) provided an indication of healthcare activity impacts on the river, and its strong correlation with ARGs indicates an additional contribution of large hospitals to the ARG input from major urban centres (**Fig. 2, 6**). The overall strong correlations of ARG concentrations in river water with each other and with other indicators of anthropogenic impact (**Fig. 6**) support ARGs as markers of anthropogenic contamination, reflecting the cumulative influence of sustained inputs from human activity along the river continuum. The consistency of results between the examined indicator ARG supports the validity of the qPCR approach, although it is limited in the number of ARG that can be observed compared to metagenomic analyses^33,34,58–60^ and the interpretation of ARG mobility, host identity, and risk evaluation that this approach allows, is valid and particularly useful for the goal of understanding AMR dynamics quantitatively.

While common sources of ARGs, metals, and nutrients primarily from wastewater discharge likely explains the observed correlations, more specific interactions such as selective effects of chemical wastewater contamination should be considered^61–63^. Downstream of WWTPs, elevated concentrations TDP closely mirrored ARG abundance. We observed a close correlation between abundant ARG and dissolved phosphorus concentrations (TDP, ρ > 0.7, median 0.79), contrasting with a weaker relationship of TDP with general bacterial abundance (*16S:* ρ = 0.53, TCC: ρ = 0.43). A weaker relationship was observed between ARG and TN (ρ > 0.65, median 0.70) or TP (ρ > 0.5, median 0.64). While P has previously been used as a proxy for bacterial growth^64^, it is not considered a selective agent for AMR. However, TDP increase was consistent with the observed enrichment of eutrophic phyla (*Proteobacteria*, *Bacteriodota*, *Firmicutes*, *Campylobacterota*^65^), which include clinically relevant genera at sites with high ARG prevalence. Such eutrophic phyla may more frequently be carriers of ARGs compared to oligotrophic phyla such as *Actinobacteriota*, *Planctomycetota*, and *Chloroflexi*^65^ which in turn declined under high P loads. The strong co-occurrence of ARGs and P likely reflects shared anthropogenic sources, such as wastewater, additional diffuse agricultural inputs, and nutrient-induced alterations of the microbial community, rather than a direct selection. Along the river (rkm 30 – 195), TP and ARG levels in agriculturally impacted areas show neither a WWTP-independent increase nor a decline, suggesting persistence of AMR markers or background inputs from diffuse sources (**Fig. 2a**).

Metals such as Cu and Zn may act as potential agents of selective pressure^66^ which could explain observed correlations with ARG (**Fig. 6**, **Fig. S9**). However, the measured Cu and Zn concentrations in both effluents and river water (**Fig. 2b**, **Fig. S3, Table S1**) remained below experimentally established thresholds required for co-selection of ARGs^67^. Nonetheless, based on reports of higher levels of Cu and other metals in influent and primary clarification, selection in the upstream wastewater stream cannot be excluded^68,69^. In effluents, despite low concentrations (< 10 µg/L), Cu and Zn correlated positively with ARGs (ρ ∼0.7 and ρ ≥ 0.6, respectively), with the strongest associations observed for *sul2* (Cu: ρ > 0.75), *aph(3’)-lb* and *ermF* (Zn: both ρ = 0.71). Correlations with *E. coli* marker *uidA* and bacterial marker *16S* were ∼0.1 lower, indicating stronger linkage of Cu to ARG-carrying subpopulations than to the overall microbial community. These findings align with Miao et al.,^64^ who also reported strong positive associations between heavy metals (Cu, Cr, Mn) and ARG prevalence, as have several similar studies^66,70,71^.

### 4.3 ARG contamination of the Aare compared to other European rivers

Can we expect a continuous accumulation of AMR along the course of major rivers? Our analysis of ARG abundance in Aare River showed their absence near the source, a consistent increase towards the middle and lower reaches, with pronounced regional hotspots. In contrast, while Danube headwaters were likewise less impacted, no correlation was found between ARGs and distance (rkm)^36^. The Rhine likewise did not exhibit a general increase with distance; instead, regionally restricted impacts were observed, particularly in the Hochrhein-Oberrhein section, attributed to the river’s size and buffering capacity against localized anthropogenic inputs^31^. Similar observations were made with respect to AMR risk indices in large Chinese rivers^33,34,60^. These patterns indicate that AMR contamination does not accumulate uniformly along rivers. Instead, source and sink dynamics can counterbalance each other, especially in larger river systems.

By comparing selected marker genes that were also measured in the Rhine and Danube studies, the AMR contamination in the three river systems can be compared. Even at the hotspot identified downstream of Bern (rkm 124), ARG relative abundance in the Aare (*intI1*: 1.7 x 10^-2^, *sul1*: 3 x 10^-3^ copies/16S) remains below the highest values reported in the Rhine (*intI1*: 6.9 x 10^0^, *sul1*: 1.07 x 10^-2^ copies/16S)^31^. Although different β-lactamase genes were assessed in the Rhine, the relative abundance at the Aare hotspot (*bla*_TEM_: 5.7 x 10^-5^) was comparable to *bla*_SHV_ levels reported in the Rhine (6 x 10^-5^, ^31^). In addition, β-lactamases in the Rhine occurred sporadically and were closely associated with WW inputs^31^, aligning with our observations for *bla_CTX-M-1_*.

Comparing absolute abundances, *intI1* (4.22 – 6.69 log_10_ copies/100 mL) and *sul1* (3.55 – 6.5 log_10_ copies/100 mL) were consistently detected at high levels across all Danube samples^30^. With the exception of the more pristine Aare headwaters, where these genes were below detection, we observed a comparable range (*intI1*: 3.67 – 6.03 log_10_ copies/100mL; sul1: 2.89 – 5.28 log_10_ copies/100mL) in the Aare. This comparison indicates that AMR contamination of the mid and lower reaches of the Aare is similar or slightly lower compared to larger European rivers, underscoring the substantial anthropogenic influence on the Aare.

In addition, load calculations showed that the Aare river acts as a major conveyor of accumulating resistance genes, with only lake passages interrupting (**Fig. S7**). This further emphasizes the role of the Aare in ARG dissemination within and beyond Switzerland’s borders, as 8.1 x 10^9^ to 5.8 x 10^11^ copies/sec of the measured ARG are discharged into the Rhine river, which enters Germany ∼50 km downstream of the confluence.

### 4.4 ARG Dynamics in a complex watershed

Tributaries exerted a moderate but notable effect on the ARG content of the Aare river. ARG content of Aare tributaries largely reflect the size of their catchments and associated human populations. The Emme, despite elevated ARG levels, contributed only modestly due to its relatively low discharge; the subsequent rise in ARGs downstream could be more plausibly attributed to effluent from WWTP Solothurn. In contrast, the Reuss and Limmat, both with large catchments and daily discharges >100 m^3^/sec (each almost 50% of the Aare discharge before their confluence) exerted a much stronger hydrological and microbial influence. Their combined effect was dilution of ARG levels (**Fig. 4**), while maintaining a high total ARG load (**Fig. S7**), as Aare’s discharge nearly doubled downstream of these confluences (from 270 to 515 m^3^/sec, **Fig. S2**). These observations underscore the critical role of absolute discharge volume on ARG distribution and dissemination.

Lakes, in contrast, behaved consistently as sinks, producing a pronounced “reset effect” on riverine ARG abundance, documented here for the first time. After passage through lakes, ARG concentrations decreased by up to 1.5 log units, with certain genes becoming undetectable in the lake outflow. These reductions appear further enhanced by high water residence time, with near-complete removal in Thunersee (648 days^72^) compared to partial reduction in Bielersee (58 days^72^), despite similar upstream ARG concentrations (**Fig. 4**). Notably, Wohlensee, a reservoir with a retention time of only 2.1 days^73^, substantially attenuated the high ARG concentrations entering from downstream of Bern. Earlier work by Czekalski et al. linked WWTP capacity in the catchment with ARG content in Swiss lakes^74^, showing that Swiss lakes are not insulated from upstream ARG inputs. The present study adds that lakes are simultaneously major ARG sinks. In large lakes with long water residence time, this likely reflects gradual replacement of the river microbiome by lake-adapted communities over time scales of months to years. The rapid response observed in Wohlensee suggests additional sink mechanisms acting on shorter timescale (days) such as sedimentation (of particle-bound cells or cell aggregates), predation, or UV-mediated decay of resistant communities under reduced flow conditions.

## Conclusions and outlook

Integration of genetic AMR markers with microbial and chemical parameters along the full course of the Aare River captured spatial dynamics, identify hotspots and sinks, and link resistance patterns to anthropogenic inputs. Our findings demonstrate that ARG levels are shaped by WW discharges and cumulative anthropogenic pressures along the entire river, positioning the Aare as a critical interface for potential AMR transmission. Our results further highlight the dynamic impact of lakes and tributaries on the main river. While tributaries can influence ARG levels through both inputs and dilution, lakes function as effective sinks that substantially reduce ARG concentrations. The behaviour observed in Wohlensee highlights how in-lake ARG dynamics remain poorly understood and require further investigation.

We also demonstrate that reporting absolute abundance (concentration), relative abundance, and total load (fluxes) of ARGs are each important parameters to assess the AMR situation of a river system and should be reported in such studies to improve comparability. This comprehensive dataset establishes a valuable baseline not only for Switzerland but also for comparable river systems, emphasizing the need for targeted management interventions. Collectively, these findings indicate that rivers function not only as passive recipients but also as active conduits of AMR, with implications that extend beyond national borders. Our study was based on a single sampling campaign, and we could not sample all potential inputs or tributaries. While our data likely represents near maximum AMR impact, repeated measurements are needed to assess temporal/seasonal variation and to investigate sources, including agricultural, in more detail. However, previous studies indicated that general patterns of AMR abundance in rivers are likely stable over seasonal timescales ^22,60,75^. Similarly, it would be interesting and valuable to perform more studies in other rivers, or to increase spatial resolution further or to analyze more potential contamination sources. Integrating metagenomic approaches into such analyses could provide valuable insights into the genetic context and risk potential of the resistome while integrating cultivation based monitoring focused on pathogens of clinical relevance can provide important information on infection risks and link to information from clinical contexts. In the context of ongoing and planned nationwide upgrades in WW treatment for organic micropollutant removal by 2040, the effectiveness of these measures should be evaluated also with respect to ARG reduction in receiving waters. Moving forward, studies that explicitly address transmission potential and risk assessment are essential.

## Acknowledgments

We thank the staff at the WWTPs in Interlaken, Thun, Bern, Biel, Lyss, Centravo AG at Lyss, Solothurn, Aarau, Oensingen and Baden for allowing and supporting effluent sampling at their facilities. We thank Antonia Zuber and Urs Lergster for the technical assistance with DNA extractions and qPCR (both Eawag, Switzerland). We thank Patrick Kathriner for performing TN, and TOC measurements, Marcel Walter for performing ICP-MS measurements, at the Aquatic Geochemistry lab at Eawag in Kastanienbaum and AuA-Lab (Analytik-und Ausbildungslabor) at Eawag Dübendorf for measuring anions by ion chromatography. We thank Dr. Martin Schmid and James Runnalls (both Eawag, Switzerland) for assisting with access to additional metadata from MetoSchweiz/Idaweb and Federal Office for the Environment (Bundesamt für Umwelt; BAFU) hydrological data, respectively. We thank Prof. Damien Bouffard (Eawag, Kastanienbaum, Switzerland) and Dr. Massimiliano Zappa (WSL, Switzerland) for valuable feedback and input on our discharge model for the Aare river. We thank Andreas Scheidegger (Statistics, Data Science & Modeling, Eawag, Dübendorf, Switzerland) for consultations regarding statistical analysis and Tara Behnsen (Surface Waters-Research and Management, Eawag, Kastanienbaum, Switzerland) and Rosi Siber (Statistics, Data Science & Modeling, Eawag, Dübendorf, Switzerland) for consultation regarding QGIS visualization and analysis. This work was supported by Swiss National Science Foundation grant 219903.

## Author Contributions

**Conceptualization**: DLW & HB (general study and all components except trace metals); DJJ (trace metal component). **Funding acquisition**: HB. **Investigation**: DLW; JA & NJA (trace metal component). **Methodology**: KB, SR, DLW, HB; DJJ (trace metal component). **Data curation**: DLW, HB. **Formal analysis**: DLW, HB; JA & DJJ (trace metal component). **Visualization**: DLW. **Software**: DLW, SR, HB. **Writing – original draft**: DLW. **Writing – review & editing**: all authors. **Supervision**: HB, KB; DJJ (trace metal component). **Project administration**: DLW, HB. **Resources**: HB; DJJ (trace metal component).

## Data Availability

Raw sequence reads of bacterial 16S rRNA gene amplicons are available from the NCBI Sequence Read Archive under BioProject PRJNA1375744. All other processed data required to reproduce the above findings are available in the manuscript and the supplementary tables.

## Declaration of generative AI and AI-assisted technologies in the manuscript preparation process

During the preparation of this work the author(s) used ChatGPT in order to revise and troubleshoot R scripts for data analysis and plotting. After using this tool/service, the author(s) reviewed and edited the content as needed and take(s) full responsibility for the content of the published article.

## References

1. Global burden of bacterial antimicrobial resistance 1990-2021: a systematic analysis with forecasts to 2050. Lancet (London, England) 404, 1199–1226; 10.1016/S0140-6736(24)01867-1 (2024).

2. Niegowska, M., Sanseverino, I., Navarro, A. & Lettieri, T. Knowledge gaps in the assessment of antimicrobial resistance in surface waters. FEMS microbiology ecology 97; 10.1093/femsec/fiab140 (2021).

3. Raaijmakers, J. M. & Mazzola, M. Diversity and natural functions of antibiotics produced by beneficial and plant pathogenic bacteria. Annual review of phytopathology 50, 403–424; 10.1146/annurev-phyto-081211-172908 (2012).

4. Ezebialu, C. U. et al. Screening and Characterization of Antibiotic Producing Organisms from Waste Dump Soil Sample. AiM 10, 422–433; 10.4236/aim.2020.109031 (2020).

5. Hwengwere, K. et al. Antimicrobial resistance in Antarctica: is it still a pristine environment? Microbiome 10, 71; 10.1186/s40168-022-01250-x (2022).

6. Andersson, D. I. & Hughes, D. Evolution of antibiotic resistance at non-lethal drug concentrations. Drug resistance updates: reviews and commentaries in antimicrobial and anticancer chemotherapy 15, 162–172; 10.1016/j.drup.2012.03.005 (2012).

7. Martin, M. J., Thottathil, S. E. & Newman, T. B. Antibiotics Overuse in Animal Agriculture: A Call to Action for Health Care Providers. American journal of public health 105, 2409–2410; 10.2105/AJPH.2015.302870 (2015).

8. Federal Office of Public Health, Swiss Confederation & Federal Food Safety and Veterinary Office. Swiss Antibiotic Resistance Report 2024 (2024).

9. Janssens, I., Tanghe, T. & Verstraete, W. Micropollutants: a bottleneck in sustainable wastewater treatment. Water Science and Technology 35, 13–26; 10.2166/wst.1997.0349 (1997).

10. Pruden, A., Pei, R., Storteboom, H. & Carlson, K. H. Antibiotic resistance genes as emerging contaminants: studies in northern Colorado. Environmental science & technology 40, 7445–7450; 10.1021/es060413l (2006).

11. Sarmah, A. K., Meyer, M. T. & Boxall, A. B. A. A global perspective on the use, sales, exposure pathways, occurrence, fate and effects of veterinary antibiotics (VAs) in the environment. Chemosphere 65, 725–759; 10.1016/j.chemosphere.2006.03.026 (2006).

12. Savin, M. et al. Antibiotic-resistant bacteria and antimicrobial residues in wastewater and process water from German pig slaughterhouses and their receiving municipal wastewater treatment plants. The Science of the total environment 727, 138788; 10.1016/j.scitotenv.2020.138788 (2020).

13. Bürgmann, H. et al. Water and sanitation: an essential battlefront in the war on antimicrobial resistance. FEMS microbiology ecology 94; 10.1093/femsec/fiy101 (2018).

14. Rizzo, L. et al. Urban wastewater treatment plants as hotspots for antibiotic resistant bacteria and genes spread into the environment: a review. The Science of the total environment 447, 345–360; 10.1016/j.scitotenv.2013.01.032 (2013).

15. Marano, R. B. M. et al. A global multinational survey of cefotaxime-resistant coliforms in urban wastewater treatment plants. Environment international 144, 106035; 10.1016/j.envint.2020.106035 (2020).

16. Ju, F. et al. Wastewater treatment plant resistomes are shaped by bacterial composition, genetic exchange, and upregulated expression in the effluent microbiomes. ISME J 13, 346–360; 10.1038/s41396-018-0277-8 (2019).

17. Read, D. S. et al. Dissemination and persistence of antimicrobial resistance (AMR) along the wastewater-river continuum. Water research 264, 122204; 10.1016/j.watres.2024.122204 (2024).

18. Jäger, T. et al. Reduction of Antibiotic Resistant Bacteria During Conventional and Advanced Wastewater Treatment, and the Disseminated Loads Released to the Environment. Frontiers in microbiology 9, 2599; 10.3389/fmicb.2018.02599 (2018).

19. S. Djordjevic & Branwen Morgan. A One Health genomic approach to antimicrobial resistance is essential for generating relevant data for a holistic assessment of the biggest threat to public health. Microbiology Australia (2019).

20. Lee, J., Ju, F., Beck, K. & Bürgmann, H. Differential effects of wastewater treatment plant effluents on the antibiotic resistomes of diverse river habitats. ISME J 17, 1993–2002; 10.1038/s41396-023-01506-w (2023).

21. Pruden, A., Arabi, M. & Storteboom, H. N. Correlation between upstream human activities and riverine antibiotic resistance genes. Environmental science & technology 46, 11541–11549; 10.1021/es302657r (2012).

22. Lee, J. et al. Unraveling the riverine antibiotic resistome: The downstream fate of anthropogenic inputs. Water research 197, 117050; 10.1016/j.watres.2021.117050 (2021).

23. Bundesamt für Umwelt BAFU. Abwasserfinanzierung / Abwasserfonds. Available at https://www.bafu.admin.ch/bafu/de/home/themen/wasser/fachinformationen/massnahmen-zum-schutz-der-gewaesser/abwasserreinigung/abwasserfinanzierung_abwasserfonds.html (2023).

24. Joss, A. et al. Biological degradation of pharmaceuticals in municipal wastewater treatment: proposing a classification scheme. Water Research 40, 1686–1696; 10.1016/j.watres.2006.02.014 (2006).

25. Knapp, C. W., Dolfing, J., Ehlert, P. A. I. & Graham, D. W. Evidence of increasing antibiotic resistance gene abundances in archived soils since 1940. Environmental science & technology 44, 580–587; 10.1021/es901221x (2010).

26. Berendonk, T. U. et al. Tackling antibiotic resistance: the environmental framework. Nat Rev Microbiol 13, 310–317; 10.1038/nrmicro3439 (2015).

27. Leonard, A. F. C., Zhang, L., Balfour, A. J., Garside, R. & Gaze, W. H. Human recreational exposure to antibiotic resistant bacteria in coastal bathing waters. Environment international 82, 92–100; 10.1016/j.envint.2015.02.013 (2015).

28. Xie, J. et al. Bacteria and Antibiotic Resistance Genes (ARGs) in PM2.5 from China: Implications for Human Exposure. Environmental science & technology 53, 963–972; 10.1021/acs.est.8b04630 (2019).

29. Weingartner, R. & Viviroli, D. Die Alpen – das Wasserschloss Europas, 2002.

30. Schachner-Groehs, I. et al. Linking antibiotic resistance gene patterns with advanced faecal pollution assessment and environmental key parameters along 2300 km of the Danube River. Water research 252, 121244; 10.1016/j.watres.2024.121244 (2024).

31. Paulus, G. K., Hornstra, L. M. & Medema, G. International tempo-spatial study of antibiotic resistance genes across the Rhine river using newly developed multiplex qPCR assays. The Science of the total environment 706, 135733; 10.1016/j.scitotenv.2019.135733 (2020).

32. Peng, F. et al. Urbanization drives riverine bacterial antibiotic resistome more than taxonomic community at watershed scale. Environment international 137, 105524; 10.1016/j.envint.2020.105524 (2020).

33. Jiang, C., Zhao, Z., Zhu, D., Pan, X. & Yang, Y. Rare resistome rather than core resistome exhibited higher diversity and risk along the Yangtze River. Water research 249, 120911; 10.1016/j.watres.2023.120911 (2024).

34. Gao, F.-Z. et al. Integrating global microbiome data into antibiotic resistance assessment in large rivers. Water research 250, 121030; 10.1016/j.watres.2023.121030 (2024).

35. Lee, K. et al. Mobile resistome of human gut and pathogen drives anthropogenic bloom of antibiotic resistance. Microbiome 8, 2; 10.1186/s40168-019-0774-7 (2020).

36. Neher, T. P., Ma, L., Moorman, T. B., Howe, A. C. & Soupir, M. L. Catchment-scale export of antibiotic resistance genes and bacteria from an agricultural watershed in central Iowa. PloS one 15, e0227136; 10.1371/journal.pone.0227136 (2020).

37. Bundesamt für Umwelt BAFU. Hydrologische Daten und Vorhersagen. Available at https://www.hydrodaten.admin.ch/de/aktuelle-lage (2025).

38. Nyall Dawson et al. qgis/QGIS: 3.36.1 (Zenodo, 2025).

39. Federal Office of Public Health, Swiss Confederation. SLMB. Determining the total cell count and ratios of high and low nucleic acid content cells in freshwater using flow cytometry. Available at https://www.admin.ch/gov/en/start/documentation/media-releases.msg-id-47549.html (2012).

40. Proctor, C. R. et al. Phylogenetic clustering of small low nucleic acid-content bacteria across diverse freshwater ecosystems. The ISME journal 12, 1344–1359; 10.1038/s41396-018-0070-8 (2018).

41. Stachler, E. et al. Quantitative CrAssphage PCR Assays for Human Fecal Pollution Measurement. Environ. Sci. Technol. 51, 9146–9154; 10.1021/acs.est.7b02703 (2017).

42. Chern, E. C., Siefring, S., Paar, J., Doolittle, M. & Haugland, R. A. Comparison of quantitative PCR assays for Escherichia coli targeting ribosomal RNA and single copy genes. Letters in applied microbiology 52, 298–306; 10.1111/j.1472-765X.2010.03001.x (2011).

43. Chern, E. C., Brenner, K. P., Wymer, L. & Haugland, R. A. Comparison of Fecal Indicator Bacteria Densities in Marine Recreational Waters by QPCR. *Water Expo*. Health 1, 203–214; 10.1007/s12403-009-0019-2 (2009).

44. Rathinavelu, S., Beck, K., Wälchli, D. L. & Bürgmann, H. Optimization and validation of a consolidated Set of TaqMan qPCR assays for the surveillance of clinically relevant antibiotic resistance genes in environmental matrices. MethodsX 15, 103600; 10.1016/j.mex.2025.103600 (2025).

45. Czekalski, N., Berthold, T., Caucci, S., Egli, A. & Bürgmann, H. Increased levels of multiresistant bacteria and resistance genes after wastewater treatment and their dissemination into lake geneva, Switzerland. Frontiers in microbiology 3, 106; 10.3389/fmicb.2012.00106 (2012).

46. Curry, K. D. et al. Emu: species-level microbial community profiling of full-length 16S rRNA Oxford Nanopore sequencing data. Nat Methods 19, 845–853; 10.1038/s41592-022-01520-4 (2022).

47. Janssen, D. J. et al. Biogeochemical cycling of trace elements and nutrients in ferruginous waters: Constraints from a deep oligotrophic ancient lake. Limnology & Oceanography 69, 2775–2790; 10.1002/lno.12687 (2024).

48. Federal Office for the Environmnet, Swiss Confederation. Hydrological data and forecasts: Discharge and water level. Available at https://www.hydrodaten.admin.ch/en/seen-und-fluesse/messstationen-zustand (2025).

49. R Core Team. R: A Language and Environment for Statistical Computing (R Foundation for Statistical Computing, Vienna, Austria, 2024).

50. Posit team. *RStudio: Integrated Development Environment for R.* (Posit Software, PBC, Boston, Ma, 2024).

51. Oksanen J, Simpson G, Blanchet F, Kindt R, Legendre P, Minchin P, O’Hara R, Solymos P, Stevens M. vegan: Community Ecology Package (2022).

52. Callender, E. & Rice, K. C. The Urban Environmental Gradient: Anthropogenic Influences on the Spatial and Temporal Distributions of Lead and Zinc in Sediments. Environ. Sci. Technol. 34, 232–238; 10.1021/es990380s (2000).

53. Comber, S. et al. Sources of copper into the European aquatic environment. Integrated environmental assessment and management 19, 1031–1047; 10.1002/ieam.4700 (2023).

54. Bau, M. & Dulski, P. Anthropogenic origin of positive gadolinium anomalies in river waters. Earth and Planetary Science Letters 143, 245–255; 10.1016/0012-821X(96)00127-6 (1996).

55. 55. Bundesamt für Umwelt. Jahrestabelle Abfluss Aare - Bern, Schönau, 2024. Available at https://www.hydrodaten.admin.ch/documents/Jahrestabellen/2135Q_24.pdf (2024).

56. MeteoSwiss. Radiation monitoring. Available at https://www.meteoswiss.admin.ch/climate/the-climate-of-switzerland/radiation-monitoring.html (2026).

57. Lee, J., Beck, K. & Bürgmann, H. Wastewater bypass is a major temporary point-source of antibiotic resistance genes and multi-resistance risk factors in a Swiss river. Water research 208, 117827; 10.1016/j.watres.2021.117827 (2022).

58. Allel, K. et al. Global antimicrobial-resistance drivers: an ecological country-level study at the human-animal interface. The Lancet. Planetary health 7, e291–e303; 10.1016/S2542-5196(23)00026-8 (2023).

59. Wang, Y. et al. Metagenomic analysis revealed sources, transmission, and health risk of antibiotic resistance genes in confluence of Fenhe, Weihe, and Yellow Rivers. The Science of the total environment 858, 159913; 10.1016/j.scitotenv.2022.159913 (2023).

60. Wang, J. et al. Supercarriers of antibiotic resistome in a world’s large river. Microbiome 10, 111; 10.1186/s40168-022-01294-z (2022).

61. Di Cesare, A. et al. Co-occurrence of integrase 1, antibiotic and heavy metal resistance genes in municipal wastewater treatment plants. Water research 94, 208–214; 10.1016/j.watres.2016.02.049 (2016).

62. Guo, X. et al. Behavior of antibiotic resistance genes under extremely high-level antibiotic selection pressures in pharmaceutical wastewater treatment plants. The Science of the total environment 612, 119–128; 10.1016/j.scitotenv.2017.08.229 (2018).

63. Raza, S., Shin, H., Hur, H.-G. & Unno, T. Higher abundance of core antimicrobial resistant genes in effluent from wastewater treatment plants. Water research 208, 117882; 10.1016/j.watres.2021.117882 (2022).

64. Miao, J. et al. Assessing the nonlinear association of environmental factors with antibiotic resistance genes (ARGs) in the Yangtze River Mouth, China. Sci Rep 13, 20367; 10.1038/s41598-023-45973-9 (2023).

65. Xie, G., Zhang, Y., Gong, Y., Luo, W. & Tang, X. Extreme trophic tales: deciphering bacterial diversity and potential functions in oligotrophic and hypereutrophic lakes. BMC Microbiology 24, 348; 10.1186/s12866-024-03488-x (2024).

66. Seiler, C. & Berendonk, T. U. Heavy metal driven co-selection of antibiotic resistance in soil and water bodies impacted by agriculture and aquaculture. Front. Microbiol. 3, 399; 10.3389/fmicb.2012.00399 (2012).

67. Kang, W., Zhang, Y.-J., Shi, X., He, J.-Z. & Hu, H.-W. Short-term copper exposure as a selection pressure for antibiotic resistance and metal resistance in an agricultural soil. Environmental science and pollution research international 25, 29314–29324; 10.1007/s11356-018-2978-y (2018).

68. Agoro, M. A., Adeniji, A. O., Adefisoye, M. A. & Okoh, O. O. Heavy Metals in Wastewater and Sewage Sludge from Selected Municipal Treatment Plants in Eastern Cape Province, South Africa. Water 12, 2746; 10.3390/w12102746 (2020).

69. Chen, A. et al. Proliferation of Resistance Genes in Wastewater Pipe Under Tetracycline and Cu Stress. Water environment research: a research publication of the Water Environment Federation 97, e70155; 10.1002/wer.70155 (2025).

70. Di Cesare, A., Eckert, E. & Corno, G. Co-selection of antibiotic and heavy metal resistance in freshwater bacteria. J Limnol 75; 10.4081/jlimnol.2016.1198 (2016).

71. Dickinson, A. W. et al. Heavy metal pollution and co-selection for antibiotic resistance: A microbial palaeontology approach. Environment international 132, 105117; 10.1016/j.envint.2019.105117 (2019).

72. Runnalls, J. Alplakes v2.0. Available at https://www.alplakes.eawag.ch/ (2025).

73. Albrecht, A., Reiser, R., Lück, A., Stoll, J.-M. A. & Giger, W. Radiocesium Dating of Sediments from Lakes and Reservoirs of Different Hydrological Regimes. Environ. Sci. Technol. 32, 1882–1887; 10.1021/es970946h (1998).

74. Czekalski, N., Sigdel, R., Birtel, J., Matthews, B. & Bürgmann, H. Does human activity impact the natural antibiotic resistance background? Abundance of antibiotic resistance genes in 21 Swiss lakes. Environment international 81, 45–55; 10.1016/j.envint.2015.04.005 (2015).

75. Amos, G. C. A. et al. Validated predictive modelling of the environmental resistome. ISME J 9, 1467–1476; 10.1038/ismej.2014.237 (2015).

