## Supplementary material for "Ripples of Resistance: Unveiling Antimicrobial Resistance Dynamics Along Switzerland’s Aare River"

**S1. Supplementary Material and Methods**

S1.1 Description of the study area

The Aare is Switzerland’s longest river, passes through the cantons Bern, Solothurn, and Aargau, draining approximately 43 % of the country (catchment size: 17709 km^2^). Originating in a largely untouched alpine region (**Fig. 1a**), fed by the Unteraar and Oberaar glaciers, the Aare River flows through a mountain valley with several villages before entering Brienzersee (29.81 km^2^) and shortly after Thunersee (47.74 km^2^) - two alpine lakes, shaped by the alpine climate and inflows from numerous mountain creeks, but also the cities of Interlaken and Thun. Downstream, the river traverses foothill landscapes of the Swiss Alps, transitioning into a mix of agricultural land and scattered settlements, with urban centres concentrated around Bern and Biel. Between them lies Wohlensee (3.65 km^2^), a slow-flowing hydropower reservoir built in a Molasse gorge, and Bielersee (39.51 km2), one of three large Jura lakes in the agriculturally productive Seeland region, fed by the Zihl canal and Jura mountain streams. From Thun to Solothurn, the river passes mainly agricultural areas, interspersed with small towns. East of Solothurn is the confluence with the Emme, a minor tributary by discharge but with a 900 km² catchment area, over half of which is farmland, originating in the Emmental Alps. Beyond Solothurn, agricultural land becomes less dominant as settlements become denser and more industrialized, particularly in the canton Aargau, before the Aare meets two major tributaries: The Reuss (draining central Switzerland) and the Limmat (draining the northeastern Alps), join the Aare before it discharges into the Rhine near Koblenz. At the confluence, the Aare River contributes more discharge than the upper Rhine.

S1.2 Flow Cytometry (FCM)

Measurements were performed on a NovoCyte Advanteon flow cytometer (Agilent, USA). For each measurement, 50 mL aliquots were taken from the original sample and stored at 4 °C until measurement (storing up to 3 days post sample collection). To prevent nozzle clogging, visibly turbid samples were passed through a 40 µm nylon Cell Strainer (Falcon, Thermo Fisher Scientific, USA) pre-measurement. As negative control, sterile-filtered Evian water purchased from the local store was used. The gating strategies for determination of total cell counts (TCC) and the high nucleic acid (HNA) and low nucleic acid (LNA) fractions are displayed in **S2, Fig. S1**.

S1.3 Quantitative PCR (qPCR)

qPCR assays met established quality criteria for linearity (R^2^ > 0.99) and efficiency (90 - 110%), except for tetA (R^2^ = 0.981) (**Tab. S7**). Slopes between -3.1 and -3.6 correspond to efficiencies between 90% and 110%, which were considered acceptable (**Tab. S7**). However, only minimal effects on measurement results are expected for the diluted samples, therefore results were normally evaluated. Negative and extraction controls were either negative or showed Cp values above the lowest standard (50 copies/reaction). Values above the LOQ were not extrapolated but set to zero to ensure consistency across genes. Outliers were excluded when standard deviations of triplicate Cp exceeded 0.5, which occurred only for values >LOQ.

Corresponding primer and fluorescence probe sequences for Taq Man assays (LightCycler 480 Probes Master; Roche, Switzerland) were adapted from internal protocols and Rathinavelu et al. (2025), except for *ermB*, *bla_CTX-M-1_*, and *qnrA*, which were analyzed using SYBR Green assay (LightCycler 480 SYBR Green Master; Roche, Switzerland). As standards for the genes *bla_CTX-M-1_*, *ermB*, *sul1*, *sul2*, *tetM*, *tetW*, *qnrA*, *intl1* and *16S rRNA*, plasmid harboring cloned inserts of the target genes, each carrying the respective target indicator gene, was utilized. Synthetic gBlock DNA, published by Rathinavelu et al. (2025) ^1^ served as standards for the genes *aadA*, *aph(3’)lb*, *bla_TEM_*, *ermF*, *mcr1*, *mecA*, *tetA,* and *uidA.* All sequences and cycling conditions are summarized in **Tables S3-S7**.

S1.4 Supplementary statistics

The relationship between ARG relative abundance and microbial community composition was assessed using complementary multivariate statistical approaches implemented in R (package *vegan*).

Pairwise associations between community composition and ARG profiles were evaluated using Mantel tests based on Spearman rank correlation between corresponding distance matrices. To compare the overall configuration of samples in multivariate space, Procrustes analysis was performed on principal coordinates analysis (PCoA) ordinations derived from community and ARG data. Statistical significance of the Procrustes association was evaluated using the PROTEST procedure. Associations between individual ARG variables and community structure were assessed using vector fitting (*envfit*) onto the community ordination.

Finally, distance-based redundancy analysis (dbRDA; *capscale*) was used to quantify the extent to which ARG profiles explained variation in microbial community composition. The significance of the overall model and of individual ARG variables was tested using permutation-based ANOVA, with sequential tests used to evaluate the independent contribution of each predictor.

All permutation tests were performed with 9,999 permutations. Statistical significance was assessed at α = 0.05.

S1.4 16S rRNA Gene Amplicon Sequencing

Amplicons were generated using the PCR cDNA barcoding kit SQK-PCB111.24 (Oxford Nanopore Technologies, UK). Undiluted DNA extracts served as templates for amplification of 16S rRNA gene (V3-V4 region) following a modified two-step PCR protocol.

PCR-1 (25 µL) contained 12.5 µL PCR mix, 9.5 µL nuclease-free water, 1 µL of each primers 341F and 806R (10 µM), and 1 µL template DNA (2 µL if concentration <10 ng/µL). Cycling conditions were 95 °C for 30 s, followed by 30 cycles of 95 °C for 15 s, 55 °C for 15 s, 65 °C for 30 s, and a final extension at 65 °C for 6 min, yielding ~460 bp amplicons. PCR-2 (25 µL) used 5 µL of PCR-1 product, 12.5 µL PCR mix, 6.75 µL nuclease-free water, and 0.75 µL barcoded primers (cDNA barcoding kit). Cycling conditions were 95 °C for 30 s, followed by 15 cycles of 95 °C for 15 s, 62 °C for 15 s, 65 °C for 40 s, and a final extension at 65 °C for 6 min, generating ~620 bp fragments. PCR products were purified with AMPure XP magnetic beads (1 µL per 1 µL PCR reaction), washed twice with 70% ethanol, and eluted in 10 µL Tris-HCl buffer (pH 8). Purified amplicons were quantified with a NanoDrop One spectrophotometer (Thermo Fisher Scientific, USA), diluted to 10 ng/µL, and pooled (≤ 100 ng total DNA per flow cell) for sequencing.

The primer pair 341f (5’-TTT CTG TTG GTG CTG ATA TTG CCC TAC GGG NGG CWG CAG-3’) and 806r (5’-ACT TGC CTG TCG CTC TAT CTT CGG ACT ACH VGG GTW TCT AAT-3’) targeted the V3 and V4 regions. Core primer sequences were originally published by Klindworth et al. and subsequently adapted in-house for improved coverage of wastewater microbiomes. Five samples failing initial library QC, likely due to matrix-related inhibition, were successfully resequenced after 10-fold dilution. Prior to sequencing, barcoded DNA libraries were pooled in equimolar ratios, and sequenced to a minimum depth of 30000 reads per sample, with most libraries yielding 40000 to 76000 reads in accordance with the manufacturer's quality standards. Reads with a Q value <10 were rejected. Median Q values per run varied between 13 and 15 per run.

FASTQ data was analyzed using the EMU software (version 3.4.5) in the MiniConda environment (version 3) using the SILVA database (Release 138, Ref Nr. 99). Data was imported into R (version 2024.04.2) and formatted for use with the Phyloseq package (by McMurdie et al.^2^) with a custom script and analyzed in R (see **Chapter 2.8**). Prior to analysis, non-bacterial sequences (such as mitochondria, chloroplasts, and eukaryotes) as well as unassigned reads, were removed.

S1.5 Metal, Chemical, and Nutrient Parameters

For nutrient analysis, 50 mL aliquots were collected in Falcon tubes to measure nutrients (TN, TP) and dissolved anions. Additionally, 10 mL aliquots were collected in glass vials for the analysis of TOC and acidified with 32% HCl in the field. Samples were stored at 4°C. For the measurement, using a Shimadzu (Japan) TOC-L analyzer, 100 mg/L C NPOC working standard solution, 100 mg/L N TN standard solution, and control solution NPOC/TN (QC 5 ppm) were prepared according to ISO 1484:1979 and ISO 11905-1:1998. Blank vials containing nanopure water enriched with 32% HCl were used. Samples and calibration standards for TP were prepared according to ISO 15681-1:2003 followed by measurements using flow-injection analysis Skalar San ++ complete system (Procon AG, Switzerland).

For metal trace quantification, samples were dried at 100 °C, digested with 0.25 mL sub-boiled HNO_3_ at 120 °C for ≥ 6 h, re-dried, and re-dissolved in 5 mL 0.2 M HNO_3_. Aliquots were transferred into acid-cleaned 15 mL polypropylene tubes for mass spectrometry analysis using an Agilent 8900 ICP-MS/MS instrument. Helium, H_2_, and O_2_ gases were used in a collision cell or as reaction gasses to minimize interferences Measurements were conducted in four batches (Sept. 2024 – Apr. 2025) with procedural blanks and certified reference material (SLRS-6) see Tab. S10. Calibration curves were prepared gravimetrically, and samples and standards were corrected via internal standardization using Rh. Additional details on analytical parameters and modes are presented in Janssen et al. (2024)^3^

**S2. Supplementary Figures**

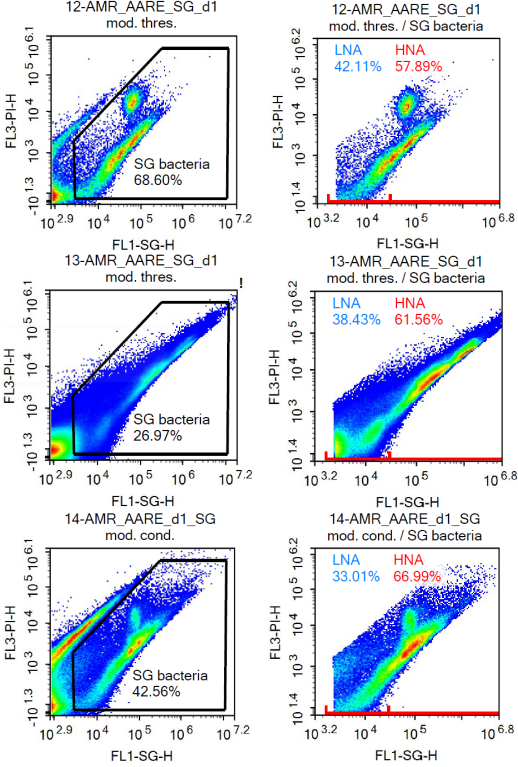

**Figure S1: Example scatter plots of the fluorescence intensity of the particles determined by flow cytometry and the gate used for the total cell count determination (SG bacteria) and high and low nucleic acid content (HNA/LNA) of three samples.** FL1-SG-H: Fluorescence intensity at the wavelength of the SYBRgreen-DNA complex. FL3-PI-H: Intensity of the nonspecific autofluorescence. Each point corresponds to a cell detection, with colors ranging from blue to red indicating accumulated signals. FCM-patterns of populations downstream of the discharge point (DS: #14) is considerably altered compared to the location upstream of the WWTP (US: #12). It shows similarities to the effluent of the treatment plant (EFF: #13).

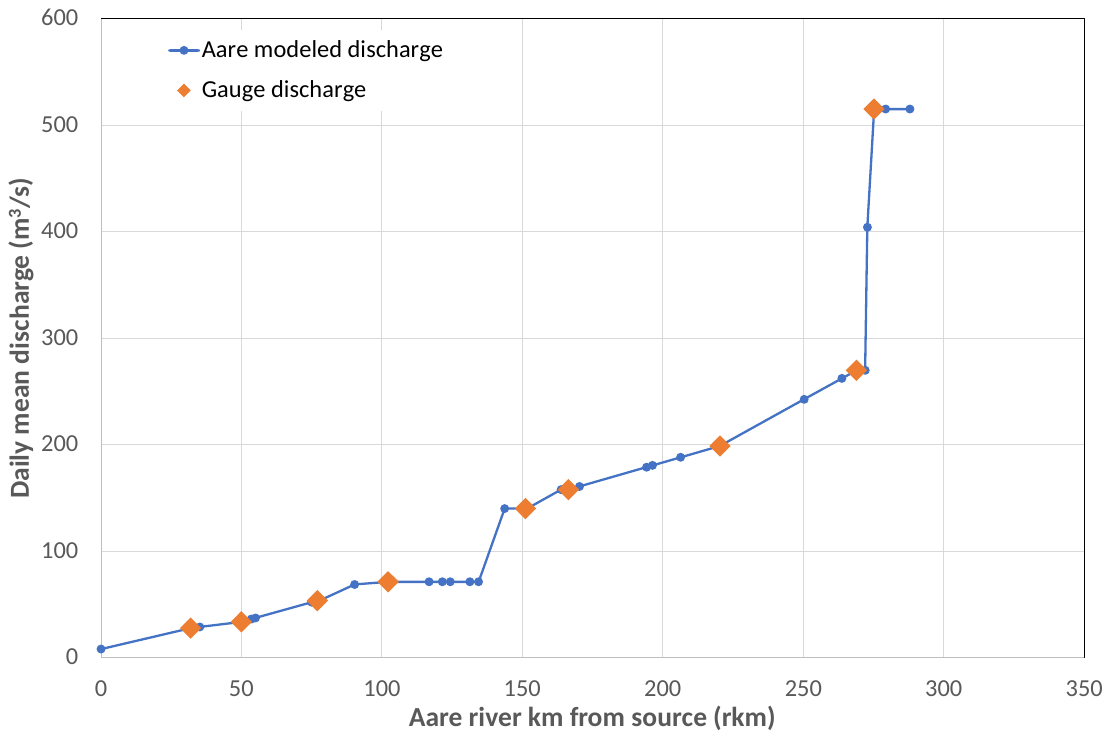

**Figure S2: Modeled mean daily discharge of the Aare river at sampling points.** Gauge discharge values (orange diamonds) were obtained for the day of sampling the nearby river reaches, see **Table S12**. Values for sampling locations (blue) were obtained by constructing a discharge model using interpolation and taking into account input from major tributaries, see **Table S13**.

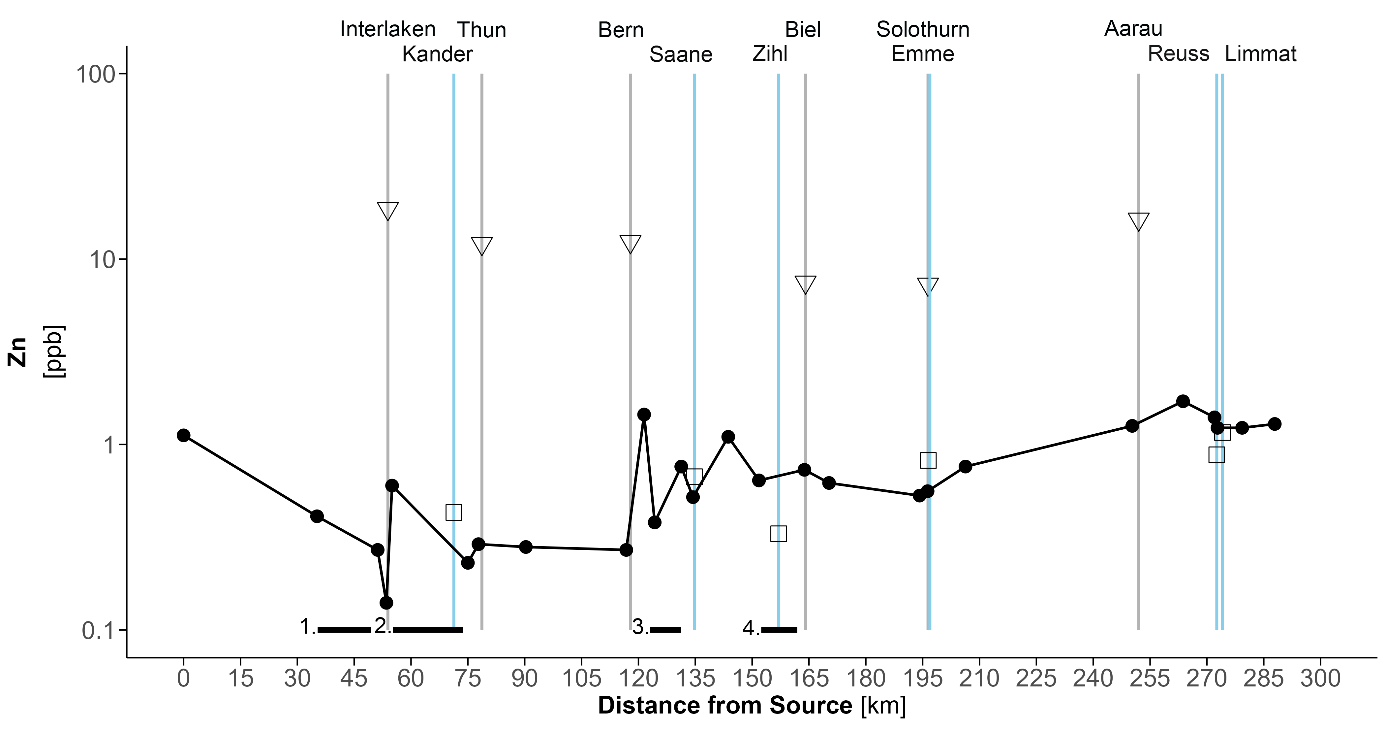

**Figure S3: Zinc (Zn) concentrations along the Aare River.** Continuous black lines represent Zn concentrations in river water along the river course. Triangles on grey bars indicate concentrations in WWTP effluents, and squares on blue bars show inputs from tributaries. Flow-through lakes are shown as black bars on the x-axis: 1) Brienzersee, 2) Thunersee, 3) Wohlensee, and 4) Bielersee. Major tributaries and corresponding WWTPs are labled on top.

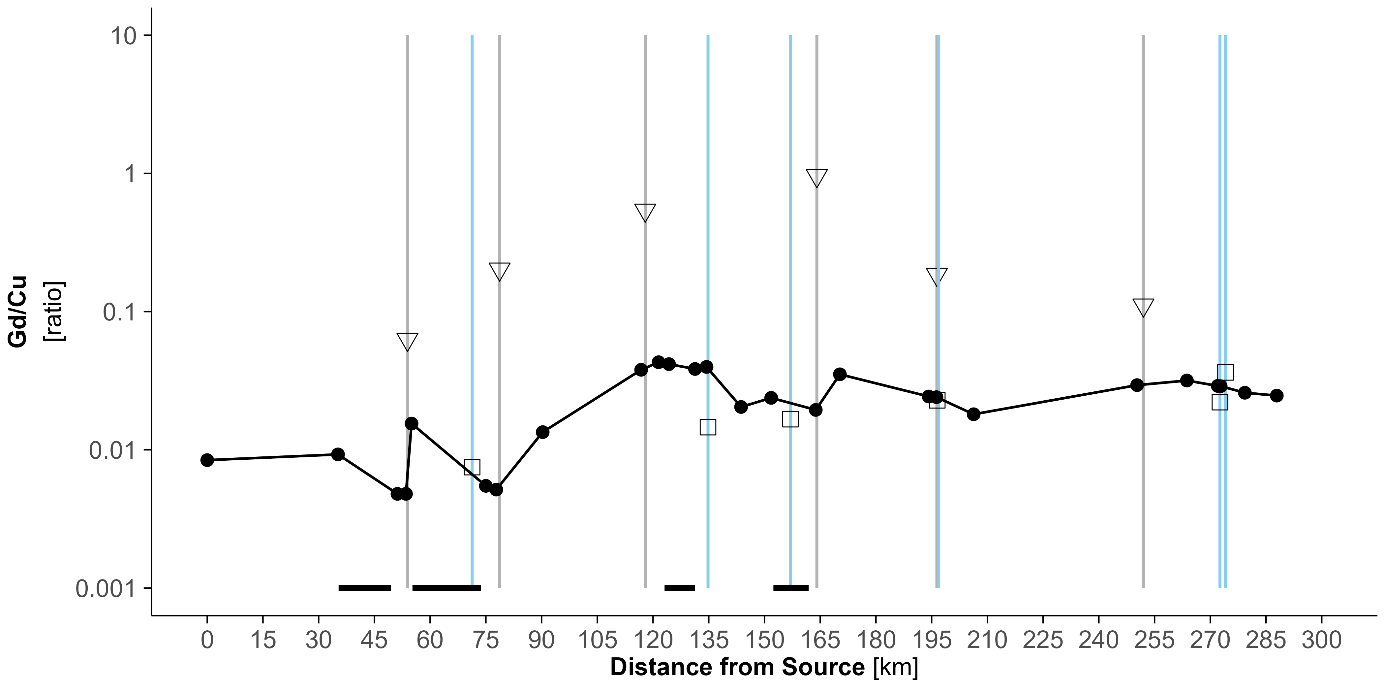

**Figure S4: Gd/Cu ratios along the Aare River.** Continuous black lines represent the ratio of Gd/Cu concentrations in river water along the river course, as an indicator of relative impact of healthcare-related activities. Triangles on grey bars indicate ratios in WWTP effluents, and squares on blue bars show ratios in the inputs from tributaries. Flow-through lakes are shown as black bars on the x-axis: 1) Brienzersee, 2) Thunersee, 3) Wohlensee, and 4) Bielersee. Major tributaries and corresponding WWTPs are labled on top.

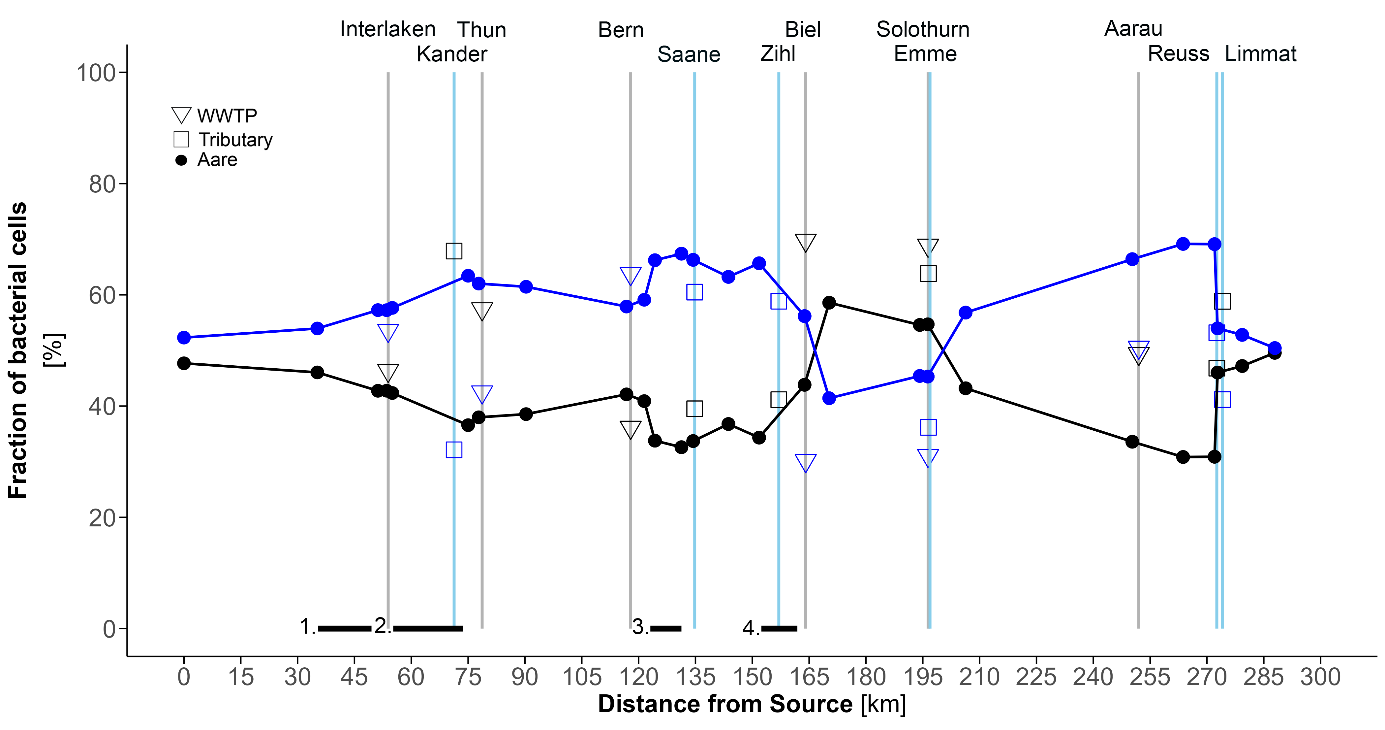

**Figure S5: Percentage of bacterial cells with low and high nucleic acid (LNA and HNA) content along the Aare River.** Fractions of LNA and HNA cells are shown along the river course (LNA: black and HNA: blue lines), in WWTP effluents (triangles on grey bars), and in tributaries (squares on blue bars). Flow-through lakes are indicated as black bars on the x-axis: 1) Brienzersee, 2) Thunersee, 3) Wohlensee, and 4) Bielersee. Raw data is provided in **Table S1**.

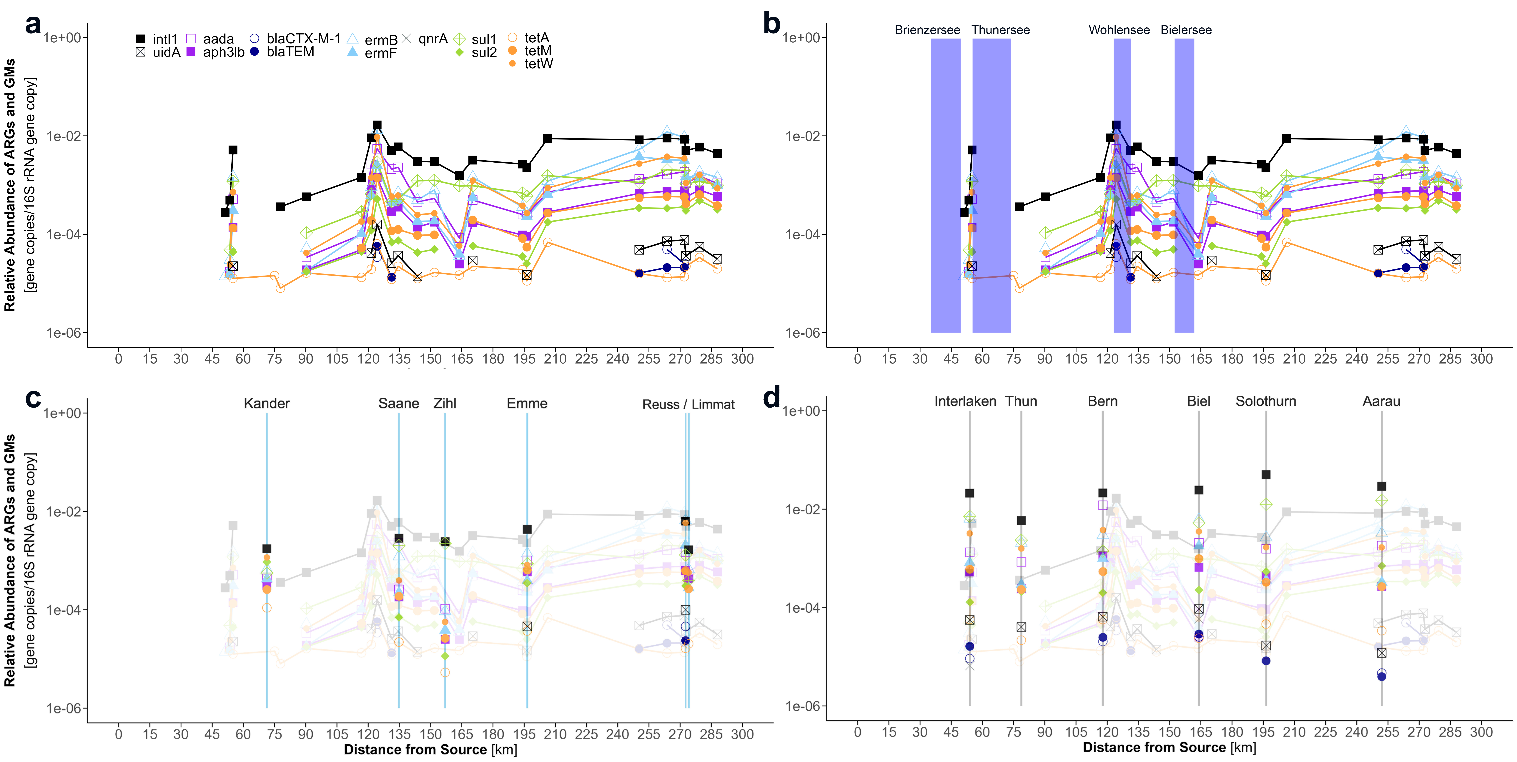

**Figure S6: Relative abundance (gene copies/16S rRNA gene copy) of ARGs and GMs, quantified using qPCR**. The concentration trends in the Aare River (a) are shown as line plots, while the abundance measured in (c) tributary inflows and (d) WWTP effluents is depicted as symbols within the bars (tributaries: blue / WWTP: gray). Flow-through lakes are indicated as bars in (b). Raw data is provided in **Table S1**.

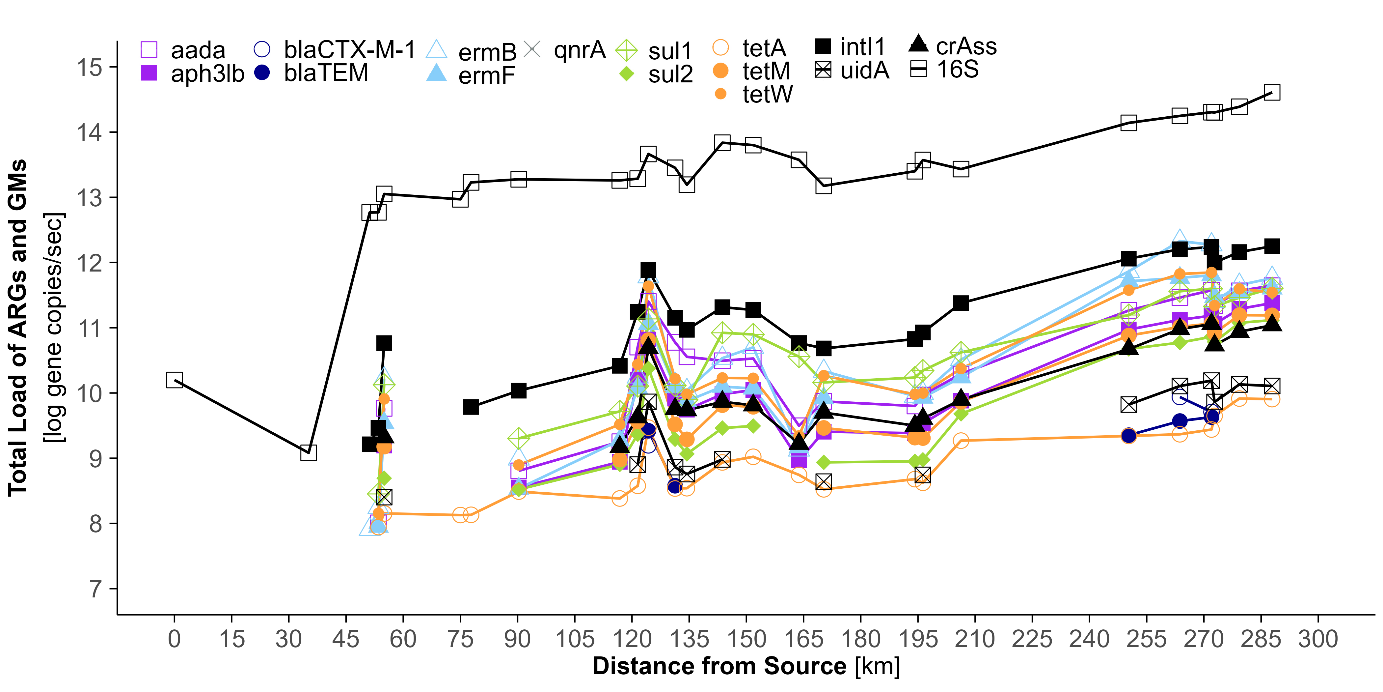

**Figure S*7*: Calculated total ARG and GM load along Aare River**. Line plots show log10-transformed mean discharge rates of ARGs and GM, normalized to river discharge. Discharge data from gauging stations along the main course and tributary inflows were used as the basis for a simple interpolation model to calculate discharge at each sampling point (see **Table S12, S13**). ARG and discharge data were then used to calculate total loads (**eq. 3**).

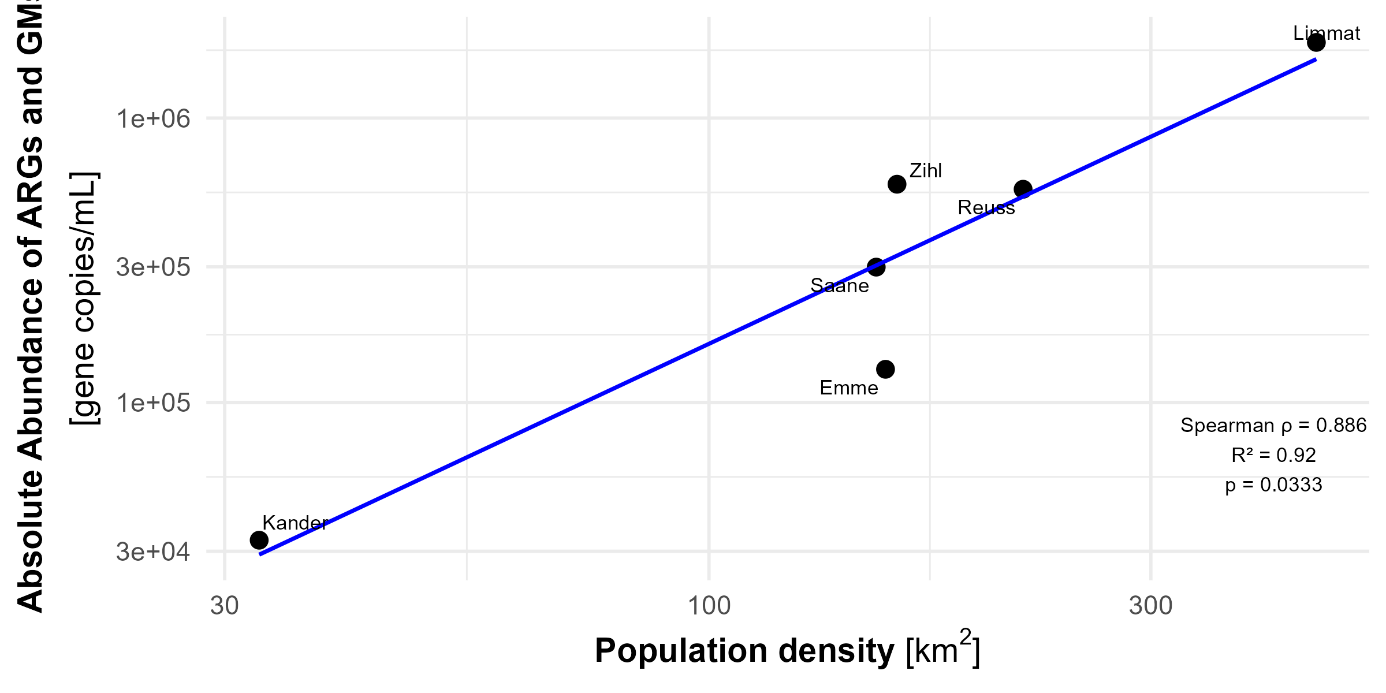

**Figure S8: Positive association between catchment population density and the summed up abundance of ARG and GM in Aare River tributaries.** Population density refers to inhabitants per km² of each tributary. The blue line shows the fitted linear regression (R^2^ = 0.92, p = 0.0333), indicating a strong correlation (Spearman ρ = 0.886) between the sum of ARG and GM abundances and population density in the tributaries' catchment.

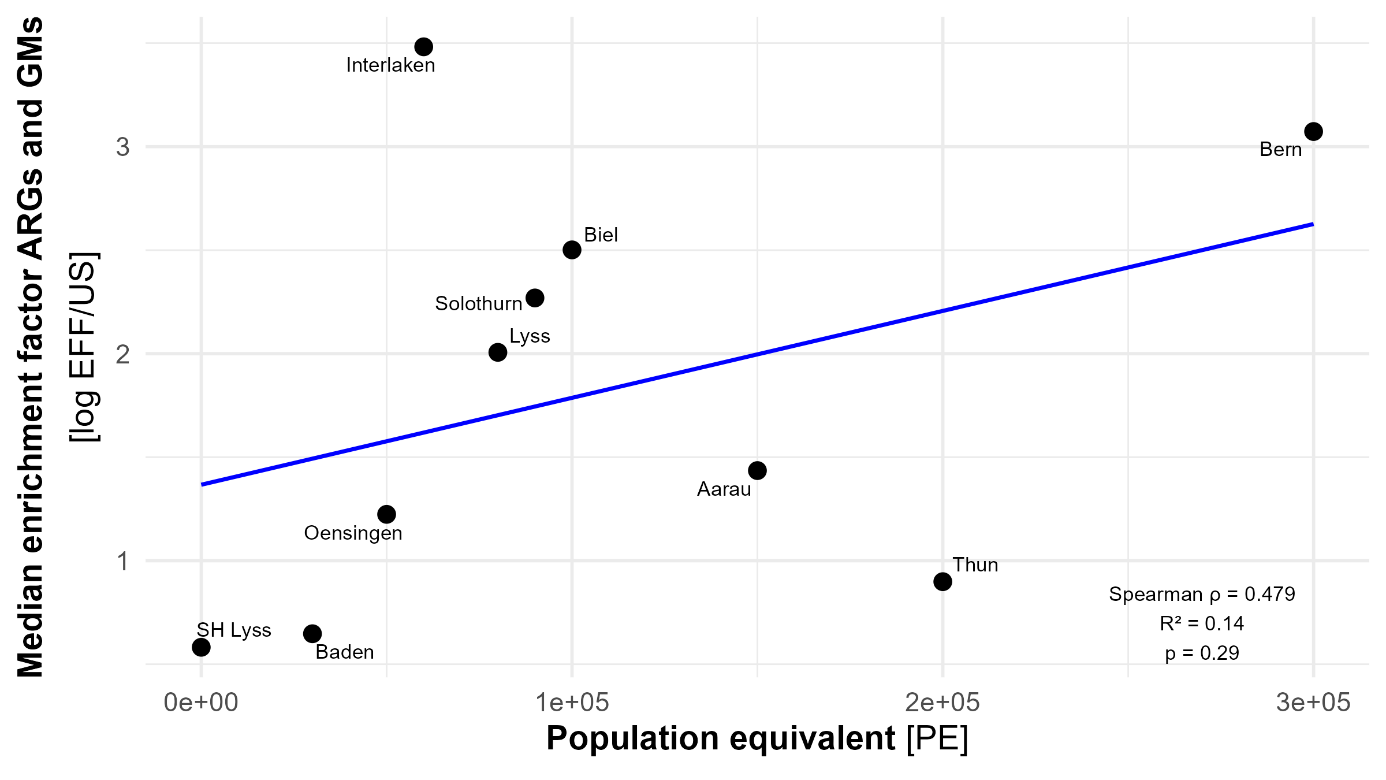

**Figure S9: Relationship between median enrichment of ARGs and GMs and design capacity of WWTPs (population equicvalents).** Each point represents one WWTP, labeled by location. The blue line indicates linear regression fit (R^2^ = 0.14), with corresponding Spearman’s rank correlation (ρ = 0.479) and p-value (p = 0.29) on the plot. A positive but non-statistical significant trend suggests slightly higher enrichment of ARGs and GMs in effluent relative to the upstream river with increasing WWTP size.

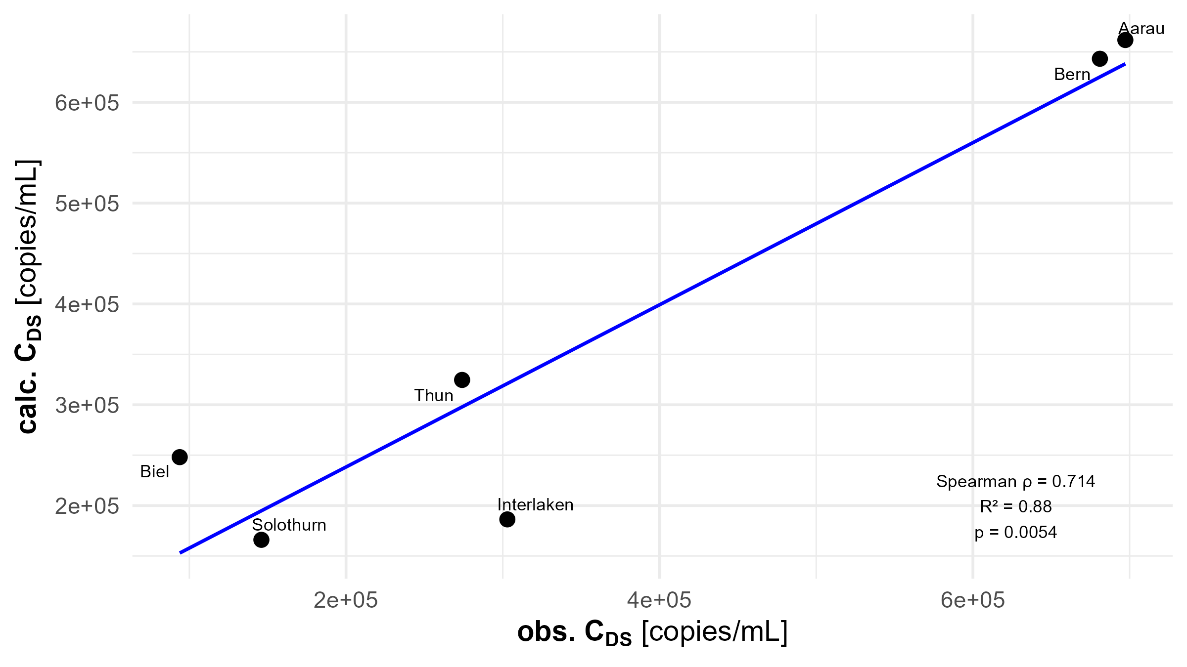

**Figure S10: Relationship between calculated and observed downstream ARG concentrations in the Aare River.** Calculated gene concentrations (calc. C_DS_) derived from the discharge-based mixing model using measured WWTP and river flow data [m^3^/s], by calculating C_DS_ = (Q_US-River_ * C_US-River_ + Q_WWTP_ * C_WWTP_) / (Q_River_ + Q_WWTP_). Observed values (obs. C_DS_) represent the sum of gene concentrations measured downstream of the WWTP effluent inlet. A strong positive association was found between calc. C_DS_ and obs. C_DS_ concentrations (Spearman ρ = 0.71, p = 0.14), and the linear regression explained most of the variance (R2 = 0.88, p = 0.005, y = 77500 + 0.804 * x). Together, these findings indicate that ARG levels downstream of WWTPs are closely linked to anthropogenic inputs from treated WW effluents.

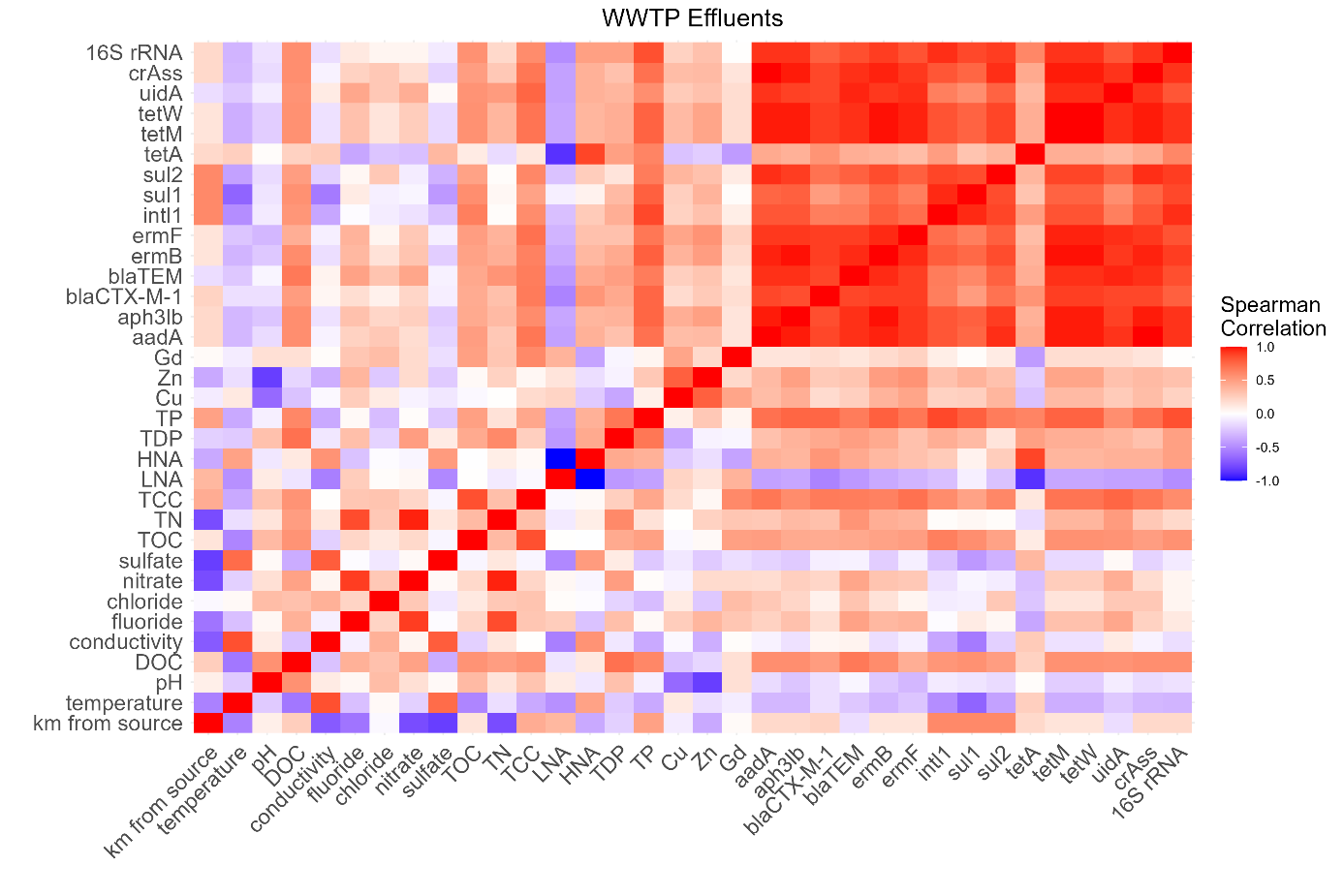

**Figure S11: Heatmap of Spearman correlation matrix, including quantified ARG and GM, physicochemical parameters, and biological indicators in all WWTP effluent samples.** Red: indicates positive correlation (ρ = 1), white: indicates no correlation (ρ = 0), blue: negative correlation (ρ= -1).

**S3 Supplementary Results**

We assessed the relationship between ARG relative abundance and microbial community composition in Aare samples using complementary multivariate analyses, including Mantel tests, Procrustes analysis, envfit, and dbRDA (see Supplementary Methods).

Across the analyses, microbial community composition and ARG profiles in Aare river samples showed a consistent but nuanced relationship. A Mantel test indicated only a weak and non-significant correlation between community and ARG dissimilarities (r = 0.16, p = 0.12), suggesting no strong global coupling. However, Procrustes analysis revealed a significant and moderate concordance between the ordination structures of both datasets (r = 0.47, p = 0.003), indicating that they share underlying gradients. Supporting this, envfit identified several ARGs (notably *sul1*, *sul2*, *intl1*, *ermF,* *aph3lb*, and *tetM*) that were significantly associated with community structure, with vectors largely aligned along a common direction in ordination space. In contrast, dbRDA showed that only a subset of ARGs (*aada, aph3lb, sul1*) explained independent variation in community composition, reflecting substantial collinearity among ARGs and suggesting that many of them act as redundant indicators of shared environmental gradients, consistent with the interpretation of wastewater inputs as the main driver. Overall, these results indicate that ARG variation is linked to microbial community structure through common ecological gradients, but this relationship is moderate and driven by a limited number of key, non-redundant ARG predictors rather than a strong overall correspondence.

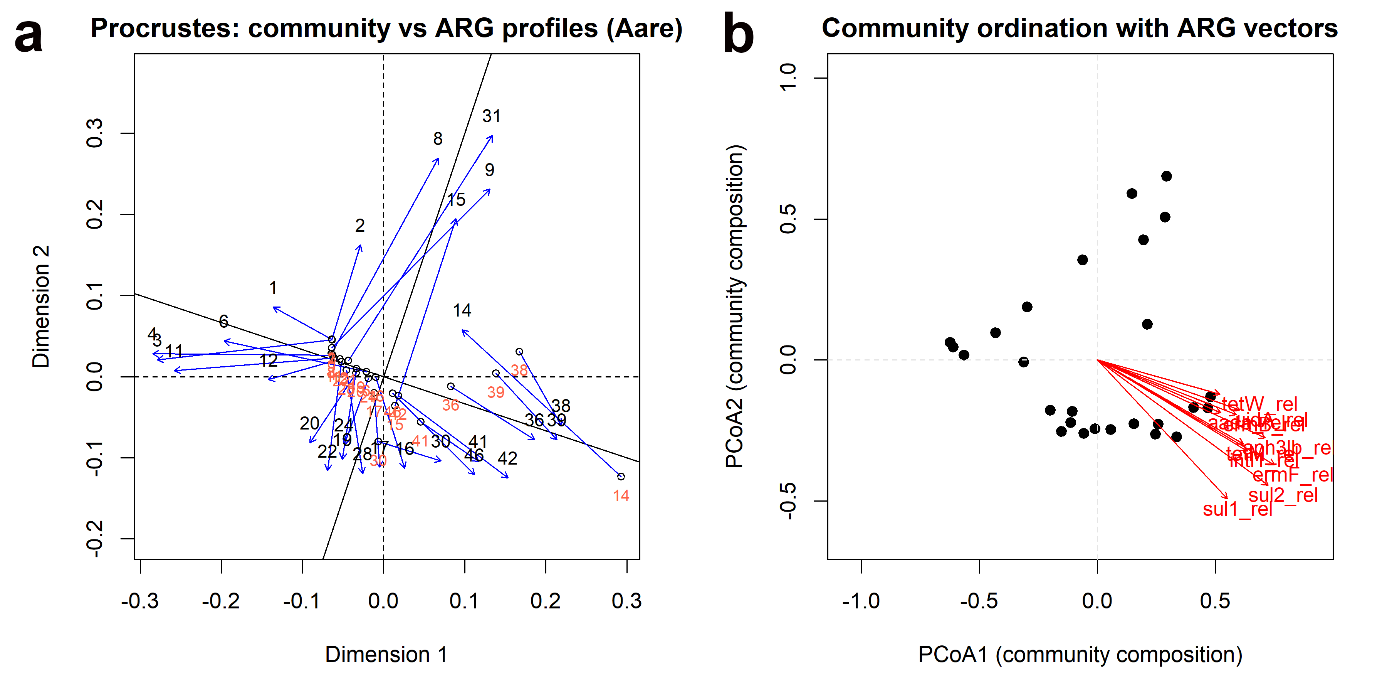
**Figure S12 relationship between ARG relative abundance and microbial community composition in Aare river samples.** (a) Procrustes analysis comparing microbial community composition and ARG profiles for Aare river samples. Points represent sample positions in ordination space, with arrows indicating the displacement between community and ARG configurations after Procrustes rotation. (b) PCoA of community composition with significant ARG vectors (envfit, p < 0.05). Arrow direction and length indicate the direction and strength of association.

**References**

1.	Rathinavelu, S., Beck, K., Wälchli, D. L. & Bürgmann, H. Optimization and validation of a consolidated Set of TaqMan qPCR assays for the surveillance of clinically relevant antibiotic resistance genes in environmental matrices. *MethodsX* **15,** 103600; 10.1016/j.mex.2025.103600 (2025).

2.	McMurdie, P. J. & Holmes, S. phyloseq: an R package for reproducible interactive analysis and graphics of microbiome census data. *PloS one* **8,** e61217; 10.1371/journal.pone.0061217 (2013).

3.	Janssen, D. J. *et al.* Biogeochemical cycling of trace elements and nutrients in ferruginous waters: Constraints from a deep oligotrophic ancient lake. *Limnology & Oceanography* **69,** 2775–2790; 10.1002/lno.12687 (2024).
